# Chromosome-level genome assembly of *Rorippa aquatica* revealed its allotetraploid origin and mechanisms of heterophylly upon submergence

**DOI:** 10.1101/2022.06.06.494894

**Authors:** Tomoaki Sakamoto, Shuka Ikematsu, Hokuto Nakayama, Terezie Mandáková, Gholamreza Gohari, Takuya Sakamoto, Gaojie Li, Hongwei Hou, Sachihiro Matsunaga, Martin A. Lysak, Seisuke Kimura

**Author notes:** **Corresponding author** Seisuke Kimura. **Competing Interest Statement:** The authors declare no competing interest.

## Abstract

The ability to respond to environmental variability is essential for living systems, especially to sessile organisms such as plants. The amphibious plant *Rorippa aquatica* exhibits a drastic type of phenotypic plasticity known as heterophylly, a phenomenon where leaf form is altered in response to the surrounding environment. Although heterophylly has been studied in various plant species, its molecular mechanism has not been fully elucidated. To establish the genetic basis and analyze the evolutionary processes responsible for heterophylly, we assembled the chromosome-level genome of *R. aquatica* by combining data from Illumina short-read sequencing, PacBio long-read sequencing, and High-throughput Chromosome Conformation Capture (Hi-C) sequencing technologies. Fine-scale comparative chromosome painting and chromosomal genomics revealed that allopolyploidization and subsequent post-polyploid descending dysploidy occurred during *R. aquatica* speciation. The genomic information above was the basis for the transcriptome analyses to examine the mechanisms involved in heterophylly, especially in response to the submerged condition, which uncovered that the ethylene and blue light signaling pathways participate in regulating heterophylly under submerged conditions. The assembled *R. aquatica* reference genome provides novel insights into the molecular mechanisms and evolution of heterophylly.

## Introduction

Plants are not able to move from their location once settled; consequently, phenotypic plasticity facilitates adaptation to fluctuating environments in the permanent habitats. One of the most striking examples of phenotypic plasticity in plants is heterophylly. Heterophylly refers to alteration of leaf form in response to environmental conditions, such as light intensity and quality, ambient temperature, and water availability (1, 2). Elucidating the mechanisms underlying heterophylly would provide insights into the strategies of adaptation of plants to fluctuating environments.

Heterophylly is often observed in amphibious plants, in which the submerged leaves are more dissected or thinner than terrestrial leaves (2). For example, submergence leads to thinner leaves in *Rorippa aquatica* (tribe Cardamineae, Brassicaceae) (3), *Hygrophila difformis* (Acanthaceae) (4, 5), *Ranunculus trichophyllus* (Ranunculaceae) (6), and *Callitriche palustris* (Callitricheae) (7). Heterophylly exhibited by amphibious plants is thought to have resulted from adaptation to fluctuating environments, particularly water level change. The evolution of heterophylly is a typical example of convergent evolution, as it has occurred multiple times independently in various taxa. Interestingly, heterophylly exhibits some similarities even at the molecular level. Previous studies have reported that ethylene is used to signal submergence. Inhibiting ethylene signaling caused leaves underwater to be similar to aerial leaves, and exogenous ethylene treatment caused thinner leaves, even under terrestrial conditions (4, 6, 7). In addition, regulation of leaf adaxial–abaxial polarity might be involved in heterophyllous leaf shape alternation in *Ranunculus trichophyllus* (6) and *Callitriche palustris* (7).

*Rorippa aquatica*, an amphibious plant found in the North American bays of lakes, ponds, and streams (8), exhibits dramatic heterophylly in response to various environmental signals, such as temperature, light quantity, and submergence (3). Its leaves become more deeply dissected and thinner underwater than in air (Fig. 1A, B). A previous study demonstrated that the mechanism of heterophylly in response to temperature in *R. aquatica*. Changes in the expression of *KNOTTED1-LIKE HOMEOBOX* (*KNOX1*) gene in response to temperature and light intensity, lead to altered concentration of gibberellins and cytokinin in the leaf primordia, which, in turn, alters leaf morphology (3).

**Figure 1.**
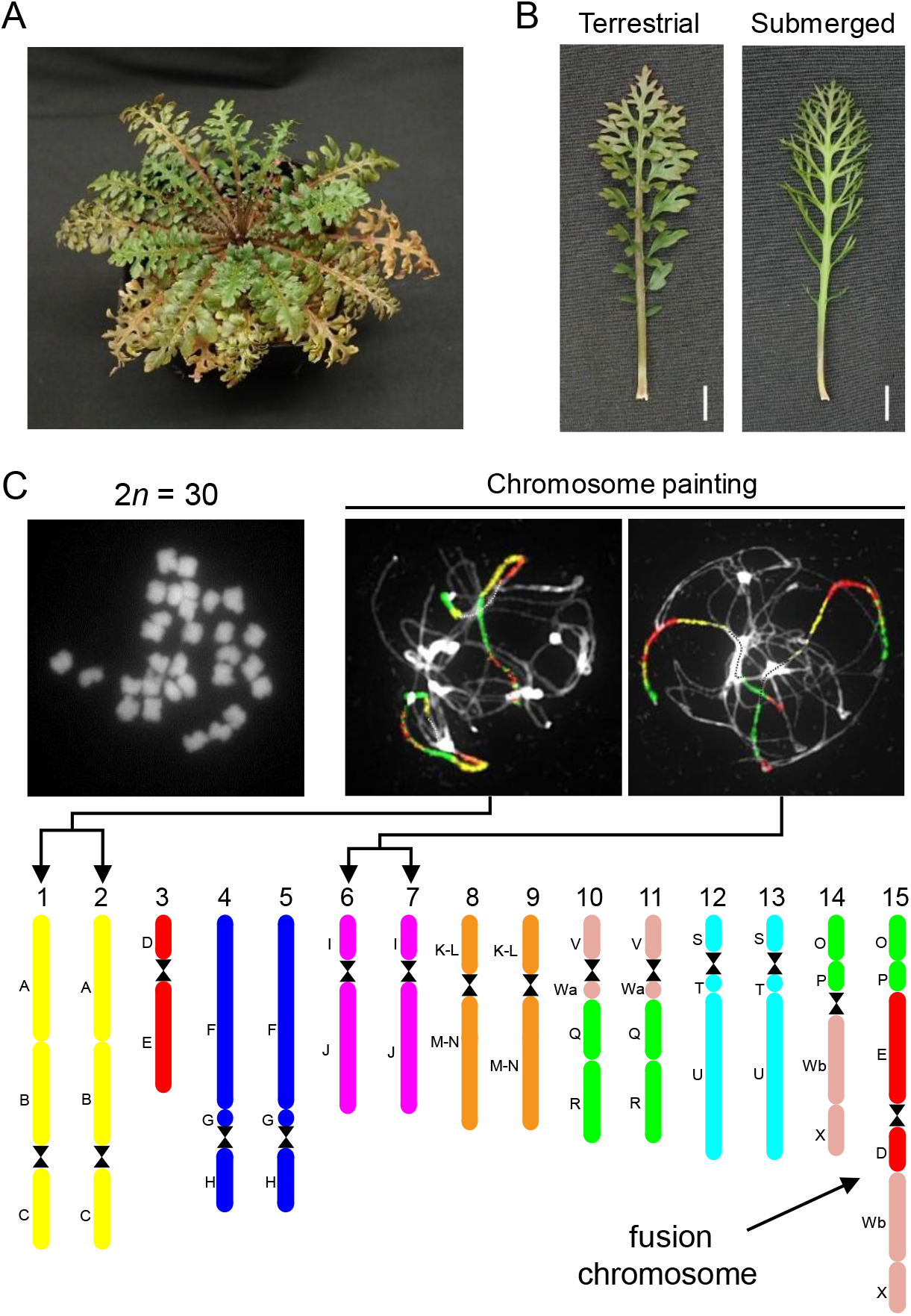
Physiological characteristics and chromosome structure of *R. aquatica*. (A) *R. aquatica* grown under terrestrial condition at 25 °C. (B) Expanded leaves of *R. aquatica* grown under terrestrial and submerged conditions at 25 °C (scale bars, 1 cm). (C) Chromosome structure of *R. aquatica*. DAPI-stained mitotic chromosomes prepared from anthers (upper left panel). Chromosome structure (lower panel) was revealed via comparative chromosome painting (Panel C, upper right). The different colors in chromosome structure correspond to the ancestral Cardamineae chromosomes, whereas capital letters refer to genomic blocks. See Fig. 2 for detailed structure of the fusion chromosome.

There have been notable advances in our understanding of the mechanisms of regulation of heterophylly in various species (3, 4, 6, 7, 9). Nonetheless, considering plants that show remarkable heterophylly are not model plants, the genomic information that could facilitate the elucidation of the underlying molecular mechanisms and evolutionary processes is lacking. *R. aquatica* is closely related to the model plant *Arabidopsis thaliana*, and *Cardamine hirsuta* (tribe Cardamineae, Brassicaceae), a model plant for compound leaf development (10–12). Therefore *R. aquatica* is potential excellent model species for studying the mechanistic basis and evolution of heterophylly. We have previously reported that the somatic cells of *R. aquatica* have 30 chromosomes (13), whereas the base chromosome number (x) in the related Cardamineae species is eight, and diploid species have 2*n* = 2x = 16 (14), which suggests that *R. aquatica* is a polyploid. In addition, the fact that the chromosome number of *R. aquatica* is not a multiple of eight suggests that its genome was restructured after polyploidization.

The evolution of plasticity is attracting considerable attention among researchers (15, 16), and understanding the evolution of heterophylly in *R. aquatica* could facilitate the elucidation of the evolutionary acquisition of phenotypic plasticity. In the present study, we performed chromosome-level genome assembly of *R. aquatica* and revealed its chromosome architecture and the evolution of its genome structure. This is the first case in which genomic information has been completed at chromosome level for a plant that exhibits remarkable heterophylly. We also combined the transcriptome data with genomic data and physiological analyses to reveal the mechanisms of heterophylly in response to submergence, and the results suggested that the response to submergence is modulated by ethylene and light signaling pathways. Our results could shed light on the molecular pathways via which heterophylly facilitates adaptation to environmental fluctuations.

## Results

### Chromosome architecture of *Rorippa aquatica* revealed via comparative cytogenomics

*R. aquatica* has 15 chromosome pairs (2*n* = 30, hereafter listed as chromosomes RaChr01 to RaChr15) (13) (Fig. 1C). We examined the *R. aquatica* genome structure and evolutionary processes via comparative chromosome painting (CCP) based on the localization of contigs of chromosome-specific Bacterial Artificial Chromosome (BAC) clones of *A. thaliana* on meiotic (pachytene) chromosomes (see Fig. 1C for examples of CCP). The painting probes were designed to reflect the system of 22 ancestral genomic blocks (GBs, labeled as A to X) (17, 18) and eight chromosomes of the ancestral Cardamineae genome (19). As all 22 GBs were found to be duplicated within the *R. aquatica* haploid chromosome complement, the species has a tetraploid origin (Fig. 1C).

We used CCP to reconstruct a complete comparative cytogenetic map of *R. aquatica* (Fig. 1C), and compared it to the ancestral genome of the tribe Cardamineae with eight chromosomes (19). Fourteen out of the 15 chromosome pairs in *R. aquatica* (RaChr01–RaChr14) are shared with the ancestral Cardamineae genome. Due to its polyploid origin, the *R. aquatica* genome contains six pairs of Cardamineae homeologues: AK1 (GBs A+B+C; RaChr01 and RaChr02), AK3 (F+G+H; RaChr04 and RaChr05), AK4 (I+J; RaChr06 and RaChr07), AK5 [(K–L)+(M–N); RaChr08 and RaChr09], AK6/8 (V+Wa+Q+R; RaChr10 and RaChr11), and AK7 (S+T+U; RaChr12 and RaChr13). Chromosomes RaChr03 and RaChr14 are homeologous to ancestral chromosomes AK2 (D+E) and AK8/6 (O+P+Wb+X), respectively. Chromosome RaChr15 (O+P+E+D+Wb+X) originated via nested chromosome insertion (NCI) of the AK2 homeologue into the centromere of the AK8/6 homeologue. At least three paracentric inversions post-dated the NCI event (Fig. 2). Comparative cytogenomic analysis confirmed the tetraploid origin of *R. aquatica* and allowed us to reconstruct the structure of the RaChr15 fusion chromosome.

**Figure 2.**
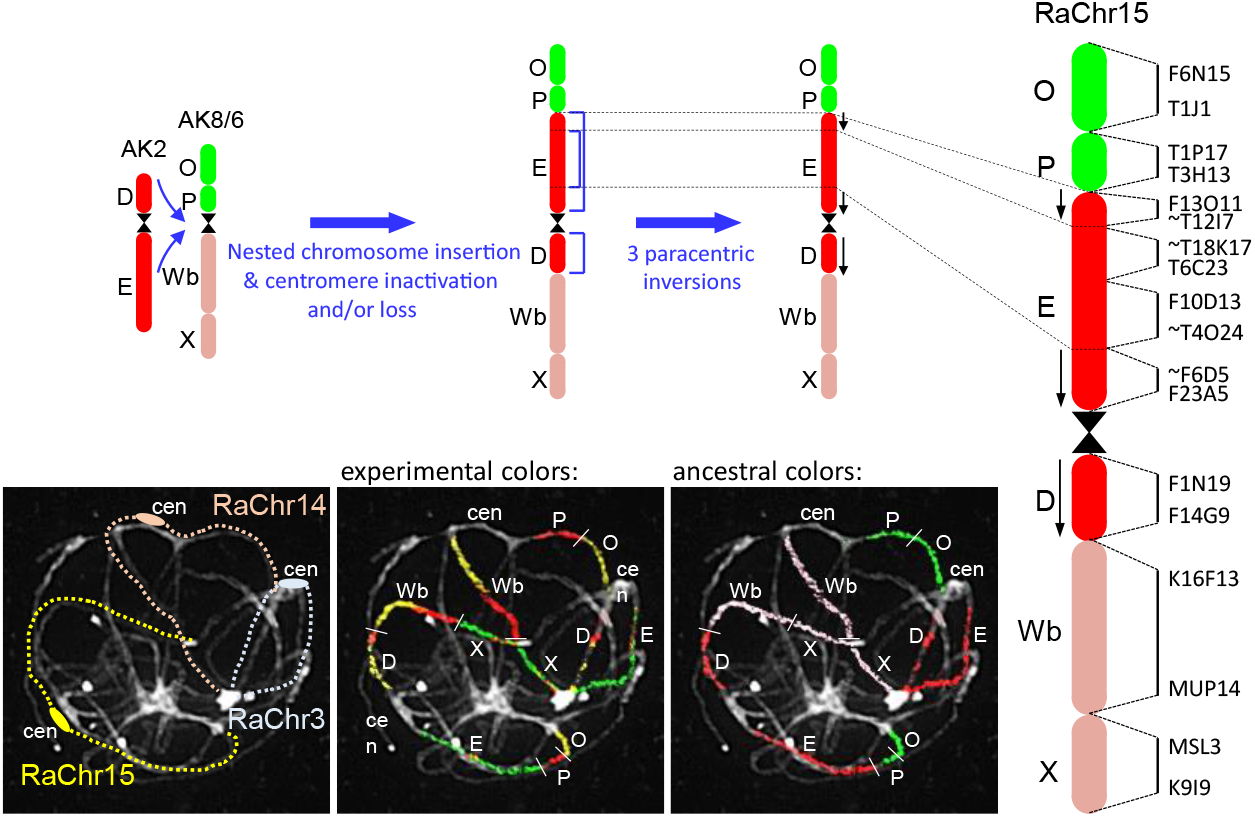
Structure of the fusion chromosome RaChr15 revealed by Comparative Chromosome Painting (CCP) of pachytene chromosomes. The different colors correspond to the ancestral Cardamineae chromosomes, whereas capital letters refer to genomic blocks. Bacterial Artificial Chromosome (BAC) clones of *Arabidopsis thaliana* defining genomic blocks or their parts are listed along the chromosome RaChr15. Centromeres are indicated by black hourglass symbols. Blue arrows and staples denote chromosome rearrangements. Black arrows along chromosome indicate the opposite orientation of chromosome regions as compared to the ancestral chromosome AK2.

### Chromosomal genome assembly and annotation

We assembled the *R. aquatica* genome and predicted its gene structure. As a reference, we used *R. aquatica* accession N, in which heterophylly was highly responsive to temperature (20). Hybrid genome assembly with Illumina short reads and PacBio long reads was performed using the MaSuRCA assembler (21). The assembled draft contigs were scaffolded using Hi-C Seq reads. Hi-C Seq provides information about the physical contact between the genomic loci in the nuclei. Therefore, given that the chromosomes are clustered in the nuclei, Hi-C Seq data scaffolding allows chromosome-level assembly. This generated 15 chromosome-level sequences and 2,040 fragments (Table S1). The total length of chromosome sequence and whole sequence were approximately 414 and 452 Mb, respectively, similar to the expected genome size (420–450 Mb) obtained using k-mer counting (Fig. S1 and Table S2). Benchmarking Universal Single-Copy Orthologs (BUSCO) was used to evaluate the assembled genome quality (Fig. S2). In total, 1,614 conserved single-copy land plant genes were screened within the *R. aquatica* genome, and 97.6% were identified, indicating the high reliability of the assembled genome. Among the 1,614 genes, 52.6% of the genes were found to be duplicated. Repeat sequences in the genome were identified (Table S3) and masked for subsequent analyses. Gene structures were predicted using the Program to Assemble Spliced Alignments (PASA) pipeline (22, 23). The results of prediction obtained using several methods were merged into an integrated gene structure, resulting in the identification of 46,197 genes (Table S4). BUSCO assessment to protein sequences from *R. aquatica* gene data identified 94.4% of the conserved genes (Fig. S2), indicating the high reliability of the gene annotation. *R. aquatica* has 1.6–1.7 times more genes than the diploid Brassicaceae species, such as *A. thaliana* (27,416 genes) and *C. hirsuta* (29,458 genes).

### Evolutionary processes revealed via comparative genomics

Although the ancestral chromosome number of Cardamineae is x = 8, *R. aquatica* has a hypotetraploid chromosome number (2*n* = 30). This chromosome number and the structure of the *R. aquatica* genome were elucidated by cytogenomic analyses (Figs. 1C and 2) as a whole-genome duplication followed by a chromosome fusion that reduced the 16 chromosome pairs to 15. To further explore this evolutionary process in *R. aquatica*, comparative genomics analysis was performed based on the chromosome sequences.

Using OrthoFinder analysis of the entire genome, a genome-level phylogeny of *R. aquatica* and related species was constructed. The longest protein sequences of each gene were extracted and used as the genome-level protein dataset in the present analysis. The datasets of 24 species were obtained from public genome databases. For intra-genus analyses, the preliminary dataset of *Rorippa islandica* (2*n* = 16) (24) was prepared by genome assembly and subsequent gene prediction using public genome-seq read data. Based on the constructed phylogenetic tree, *R. aquatica* and *R. islandica* were placed in the clade that includes *Arabidopsis* (Fig. S3), corresponding to the Brassicaceae clade A (25). *Barbarea vulgaris* is the closest species and *C. hirsuta* is the closer sister of the genus *Rorippa*.

To investigate the origin of the *R. aquatica* genome, we compared the chromosome-level assemblies of *R. aquatica* and *C. hirsuta* (Fig. S4). Multiple alignment based on nucleotide similarity showed that each chromosome of *C. hirsuta* was similar to two *R. aquatica* chromosomes, indicating a tetraploid origin of the latter species. The collinearity of the RaChr15 fusion chromosome with the *C. hirsuta* chromosomes Chr2 and Chr8 indicates that this *R. aquatica* chromosome was formed via an NCI involving the *Rorippa* homeologues RaChr03 and RaChr14, consistent with our CCP-based results (Fig. 2).

The divergence ages of duplicated genes could indicate when *R. aquatica* attained its tetraploid-like characteristics. To estimate divergence ages in the Brassicaceae, we selected eight species (*A. thaliana, A. lyrata, Barbarea vulgaris, Capsella rubella, Cardamine hirsuta, Eutrema salsugineum, R. aquatica*, and *R. islandica*). All genes in each genome were clustered into orthogroups (i.e., groups consisting of orthologous genes) based on protein similarity. In total, 10,845 single-copy orthogroups were found upon comparing six of the eight species (excluding *R. aquatica* and *R. islandica*). In *R. islandica*, almost all (91.2%) were single-copy genes, whereas in *R. aquatica*, 58.6% were duplicated genes (Table S5). This suggests that the *R. aquatica* genome underwent large-scale gene duplication. To estimate species divergence and gene duplication ages, we calculated the synonymous nucleotide substitution rate (Ks) between gene orthologs, both inter- and intra-species. To calculate Ks, we used the longest coding DNA sequences of each gene conserved as a single copy in the seven Brassicaceae species and duplicated in *R. aquatica*; this resulted in 5,856 sequences. The Ks distributions of *R. aquatica* compared with other species and those of *R. islandica* were quite similar, reflecting phylogenetic relatedness (Fig. S5). Divergence ages were estimated using Ks as T (in years) = Ks / (2 * *μ*), where *μ* = 6.51648E−09 synonymous substitutions/site/year for Brassicaceae (26). On that basis, the divergence times between *Cardamine* and *Rorippa*, and between *Barbarea* and *Rorippa* are 13.7–14.2 million years ago (Mya) and 10.5–10.8 Mya, respectively (Table S6). The Ks distribution based on the duplicated paralogs of *R. aquatica* has a single peak with a median of Ks = 0.102, corresponding to a divergence of 7.8 Mya. This suggests that large-scale gene duplication in *R. aquatica* occurred after the *Barbarea*/*Rorippa* divergence, and most likely occurred at the whole-genome level. In *R. aquatica* and *R. islandica*, the median of Ks is 0.073, corresponding to a divergence of 5.6 Mya. Although *R. islandica* has diploid characteristics (Table S5) and chromosome number of 2*n* = 16 (24), our findings indicate that these two *Rorippa* species diverged after the whole-genome duplication (WGD).

To elucidate this conflict, Ks between *R. aquatica* and *R. islandica* was analyzed at the chromosome level. The chromosomes of *R. aquatica* form two subgenome groups based on their distribution of Ks calculated relative to the *R. islandica* genome (Fig. 3A and Table S7, S8). The first group, named subgenome A, includes eight chromosomes (RaChr01, -03, -05, -07, -08, -10, -13, and -14) with a median Ks value of approximately 0.05. These chromosomes are closer to those of *R. islandica*. Subgenome B includes seven chromosomes (RaChr02, -04, -06, -09, -11, -12, and -15) with a median Ks value of approximately 0.09. These chromosomes are phylogenetically more distant from those of *R. islandica*; their Ks values relative to *R. islandica* are similar to those between the duplicated *R. aquatica* genes. The fact that *R. aquatica* has two subgenomes with different divergence ages indicates that it has an allotetraploid origin caused by hybridization between two ancestral *Rorippa* species. Integration of various genomic analyses clarified the structure of the *R. aquatica* genome (Fig. 3B). The homologous chromosomes in each subgenome show a similar distribution of genes and long terminal repeats (LTRs). The peaks of the LTR distribution in each chromosome indicate centromeric regions. These results also suggest that gene duplication in the species occurred via a WGD.

**Figure 3.**
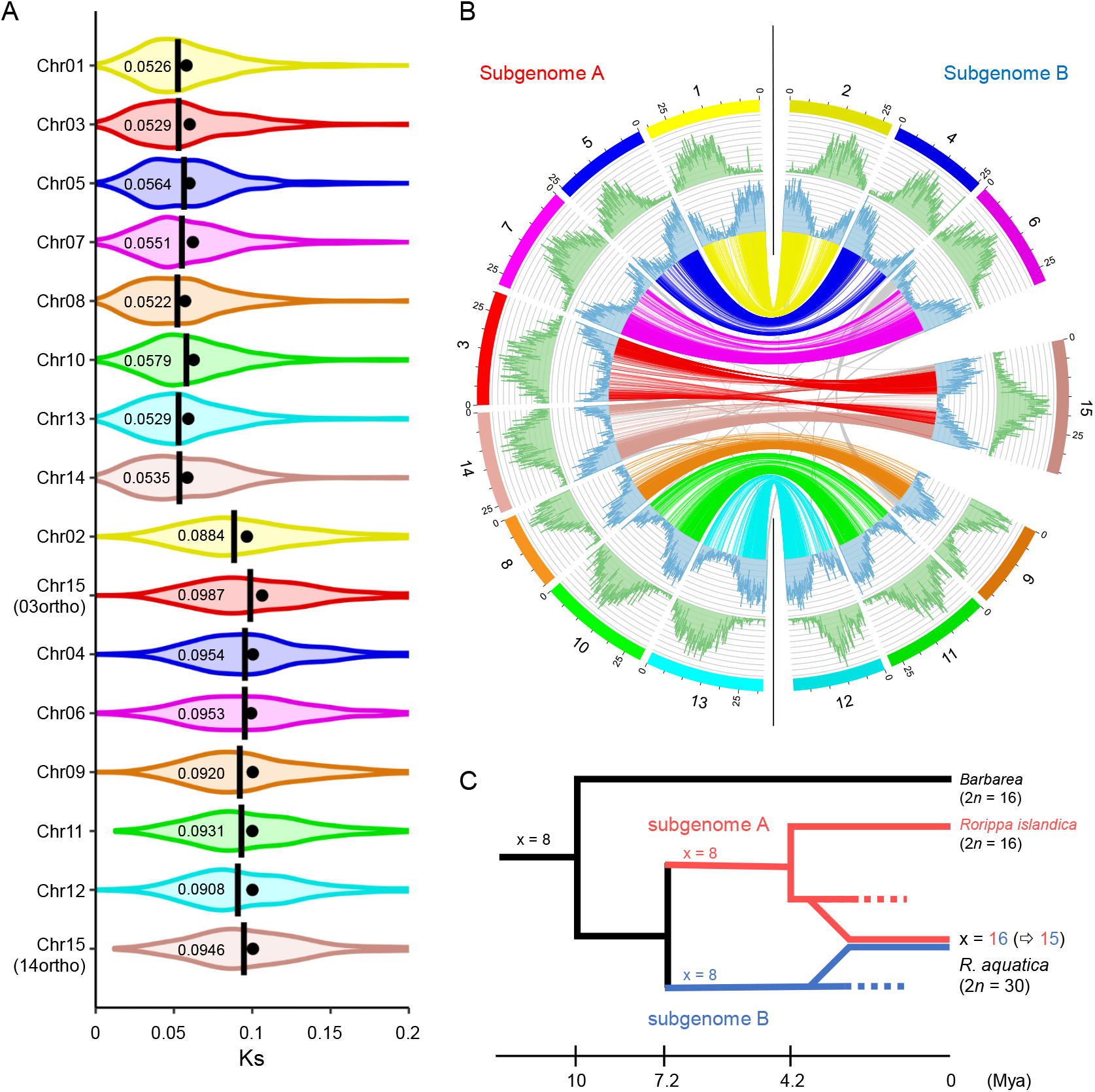
*Rorippa aquatica* chromosome-level genome assembly. (A) Chromosome-level synonymous nucleotide substitution rate (Ks) distributions relative to the *R. islandica*, for *R. aquatica* paralogs in orthologous chromosomes. Closed circles: mean; bars and numbers: median. (B) Circos plot of the assembled *R. aquatica* genome. In the Circos plot, the long terminal repeat and gene distributions are in green and blue, respectively. The lines at the center link the paralogous genes for which orthologous genes were conserved as single copy in diploid Brassicaceae species. (C) Evolutionary scheme of the formation the allotetraploid genome of *R. aquatica* based on the present data.

Comparative genomics revealed the evolutionary process of establishing the present *R. aquatica* (Fig. 3C): assuming an allotetraploid origin of *R. aquatica, Rorippa* split into the subgenome groups A and B approx. 7.2 Mya (Ks = 0.09). In the subgenome A group, divergence into the ancestor of *R. islandica* and the parental species of *R. aquatica* occurred 4.2 Mya (Ks = 0.05). Subsequent hybridization of two species from different subgenome groups resulted in the formation of the allotetraploid origin of *R. aquatica*. Phylogenetic analysis based on plastid sequences placed *R. islandica* in a clade distant from *R. aquatica* (13). This suggests a paternal origin of subgenome A of *R. aquatica*. Our analysis does not identify the seed parent of subgenome B. Based on Ks analysis, the fusion chromosome RaChr15 was formed by intra-subgenomic fusion. The median Ks value of RaChr15 versus *R. islandica* was approximately 0.09, similar to the other chromosomes of subgenome B (Fig 3A and Table S7). Chromosomes RaChr03 and RaChr14, within subgenome A, remained as independent chromosomes.

### Transcriptome analysis reveals the pathway underlying heterophylly in response to submergence

To elucidate the mechanism of heterophylly in response to submergence, we conducted RNA-seq gene expression analysis using the assembled *R. aquatica* genome data. First, we observed the morphology of young leaves over time to determine the timing of leaf shape change upon submergence (Fig. 4A). After 1 day of submergence, submerged and terrestrial leaves did not differ morphologically. After 4 days of submergence, the young-leaf margin serrations became deeper in the submerged compared to in terrestrial plants. After 7 days of submergence, leaf incisions were significantly deeper in the submerged leaves. To reveal the gene expression patterns of early response to submergence as well as the early stage of leaf morphology differentiation, shoot apices containing young leaves were sampled at 1 hour and 4 days after submergence and RNA-seq analysis was performed. As a result, we identified 787 upregulated and 1,091 downregulated genes 1 hour after submergence, which increased to 5,358 upregulated genes and 4,945 downregulated genes after 4 days of submergence (Fig. 4B). The submergence-responsive differentially expressed genes (DEGs) were classified into three classes according to the timing of expression. The genes whose expression changed only within 1 hour of submergence were classified as “early response genes.” The genes whose expression changed after 1 hour as well as after 4 days of submergence were classified as “throughout response genes.” The genes whose expression changed only after 4 days of submergence were classified as “late response genes.”

**Figure 4.**
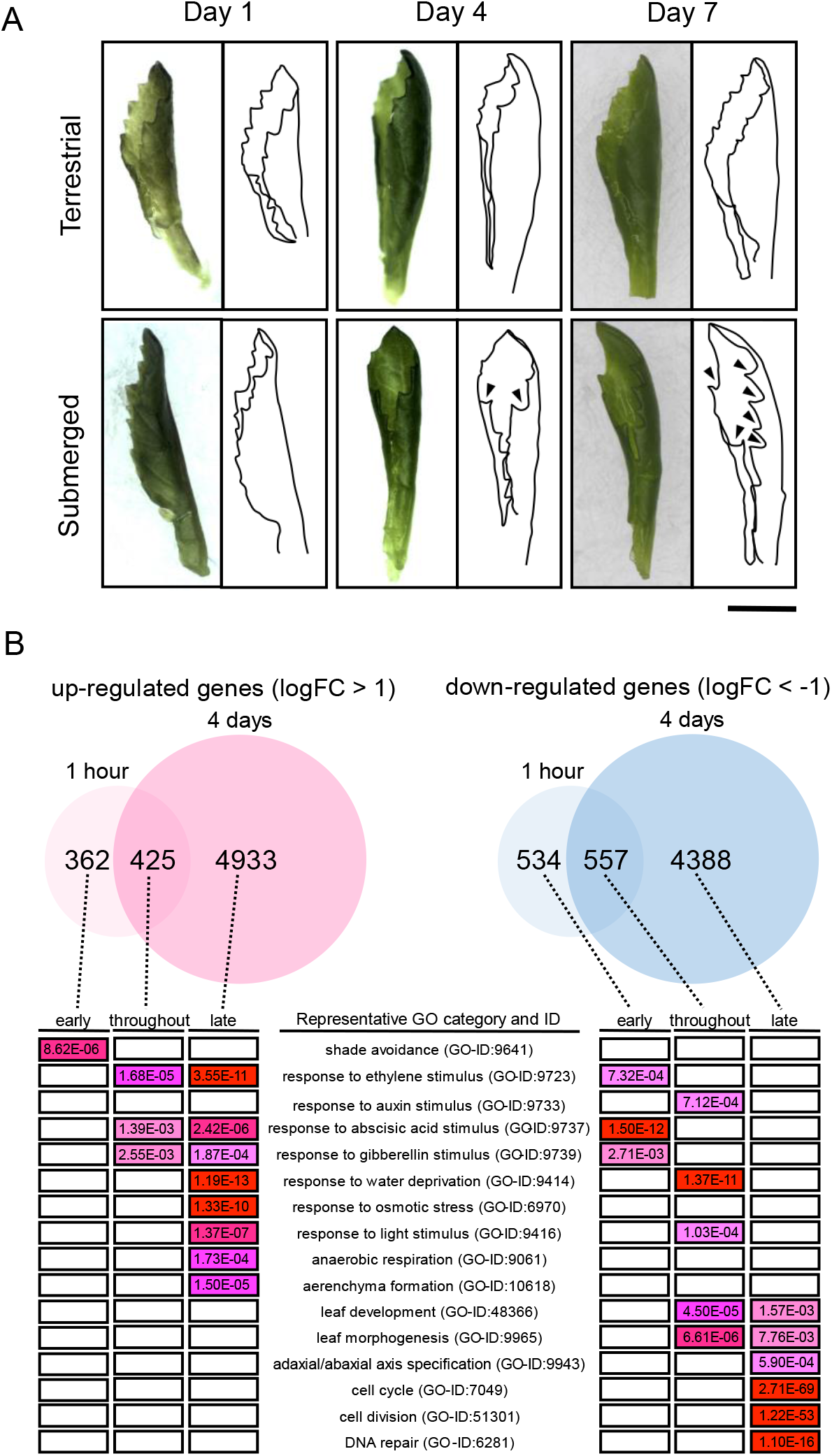
The effect of submergence on *Rorippa aquatica* leaf morphology and gene expression. (A) Images and outlines of newly emerged young leaves after transfer to terrestrial or submerged conditions (scale bar, 1 cm). (B) Transcriptome analysis of leaves grown under the submerged condition. Differentially Expressed Genes (DEGs) were identified based on significant differences in expression and log fold change |LogFC| > 1. Based on their expression patterns, DEGs were categorized as “early response genes” (responding only within the first hour), “late-response genes” (after 4 days), and “throughout-response genes” (throughout submergence), then subjected to Gene Ontology enrichment analysis (significantly enriched categories are shown).

Next, we used Gene Ontology (GO) enrichment analysis to elucidate the biological processes involved in regulating heterophylly (Fig. 4B and Supplementary information 1). Among early response upregulated genes, those in the “shade avoidance” category (GO-ID: 9641) were enriched, suggesting that light conditions are important in the early response to submergence. Significant numbers of both up- and down-regulated genes were related to phytohormones, such as ethylene (GO-ID: 9723), gibberellin (GO-ID: 9739), and abscisic acid (GO-ID: 9737). Some of the down-regulated genes were enriched in “response to auxin stimulus” (GO-ID: 9641). The results are consistent with previous findings (20) that phytohormones participate in regulation of heterophylly. Genes related to aspects of leaf morphology such as leaf development (GO-ID: 48366) and leaf morphogenesis (GO-ID: 9965) were downregulated in response to submergence. Those involved in adaxial/abaxial axis specification (GO-ID: 9943) were downregulated during the late response. Regulation of cell division and elongation is essential in altering leaf morphology. At the late stage, when the leaf morphology differed between the submerged and terrestrial leaves, genes belonging to the cell cycle (GO-ID: 7049) and cell division (GO-ID: 51301) categories were enriched among the downregulated genes. The results indicate that the expression of genes involved in leaf development is regulated immediately after submergence, and that phytohormones are involved in the regulation.

### Ethylene induces the submerged-leaf phenotype

The gene expression profiling revealed the importance of gibberellin in *R. aquatica* heterophylly, which is consistent with the previous study (3). Furthermore, abscisic acid is vital for the response to aquatic environments (4, 7, 27). Nonetheless, the relationship between ethylene and heterophylly in *R. aquatica* remains to be elucidated.

In numerous plant species, ethylene accumulation in plant tissue during submergence triggers the submergence response; for instance, ethylene participates in heterophylly in some plant species (28). Therefore, we examined the relationship between ethylene and *R. aquatica* leaf shape. Treating terrestrial *R. aquatica* plants with 1-aminocyclopropane-1-carboxylic acid (ACC), an ethylene precursor, resulted in the formation of more deeply lobed leaves, with thinner leaf blades than those in the untreated terrestrial plants (Fig. 5A). In contrast, when the ethylene-response inhibitor AgNO_3_ was added under submerged conditions, leaves with expanded blades, similar to the terrestrial leaves, were formed (Fig. 5B). Further, as the concentration of ethylene was increased, thinner leaf blades developed (Fig. 5C). The results indicate that heterophylly in *R. aquatica* is regulated by the levels of ethylene hormone. The submerged-phenotype of leaf was suppressed by inhibiting the ethylene response even under submerged conditions, suggesting that the regulation of heterophylly is mediated by the ethylene response pathway.

**Figure 5.**
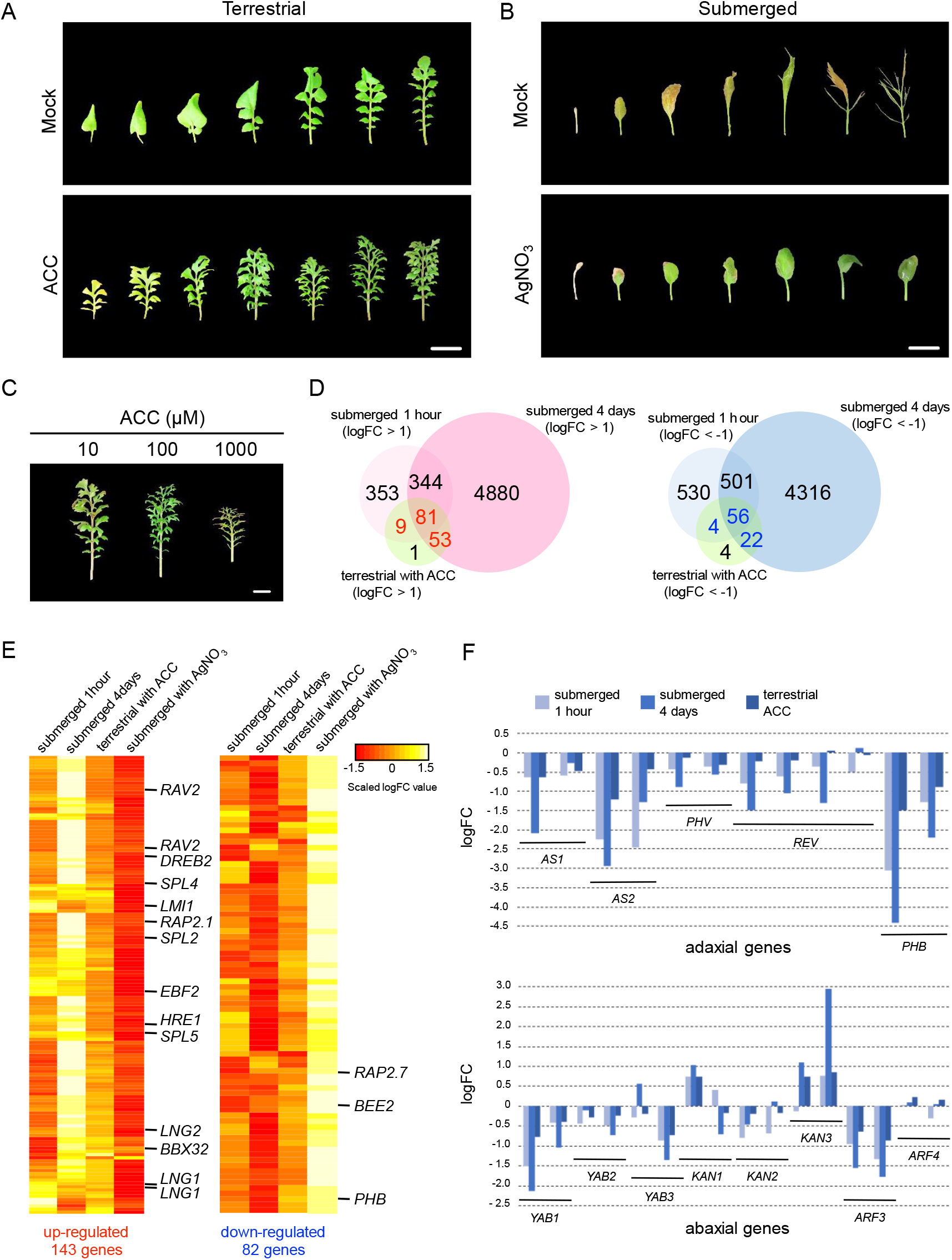
Effect of ethylene on *Rorippa aquatica* heterophylly. Leaves produced after treatment with (A) mock or 100 µM 1-aminocyclopropane-1-carboxylic acid (ACC, an ethylene precursor) under terrestrial conditions, or (B) mock or 1 µM AgNO_3_ (ethylene inhibitor) under submerged conditions (scale bars, 1 cm). (C) Ethylene-dose effect under terrestrial conditions, with different ACC concentrations (scale bars, 1 cm). (D) Selection of candidate genes that induce submerged-phenotype leaves: genes that were up- or down-regulated under either submergence or ethylene treatment were used as candidates. (E) Candidate gene expression profiles. (F) Responses of adaxial–abaxial polarity determining genes to the submergence and ethylene treatments.

Next, we performed RNA-seq analysis of ACC-treated plants to identify the genes responsible for submerged-type leaf formation. Since ACC treatment induced the formation of submerged-type leaves, we extracted genes whose expression changed both under submerged and ACC-treated conditions (Fig. 5D, E and Supplementary information 2): 143 genes were commonly upregulated, and 82 genes were commonly down-regulated. As expected, several common ethylene response genes were upregulated in both datasets.

One of the commonly regulated genes, *LONGIFOLIA* (*LNG1* and *LNG2*), has been identified to participate in leaf morphogenesis in *A. thaliana* via activation-tag gene screening (29): the dominant mutant of *LNG1* (*lng1-1D*) formed elongated and narrow leaves with serrated margins; furthermore, *LNG1* and *LNG2* may work redundantly and regulate longitudinal cell elongation.

*LATE MERISTEM IDENTITY1* (*LMI1)* and *REDUCED COMPLEXITY* (*RCO*) arose from the same ancestral gene via gene duplication within a clade of Brassicaceae (30). *RCO* controls leaf complexity (30) in *C. hirsuta* wherein the wildtype has compound leaves but the *rco* mutant displays simple lobed leaves. *RCO* is lost and *LMI1* remains in *A. thaliana* emerging simple leaves, and the introduction of *C. hirsuta RCO* into *A. thaliana* resulted in the formation of serrations. The *R. aquatica* gene *RaChr03G09000*, extracted as an *A. thaliana LMI1*, is an *RCO* ortholog, because of higher protein identity to *C. hirsuta RCO* (84.3%) than *C. hirsuta LMI1* (63.1%). The upregulation of *R. aquatica RCO* might cause compound leaf formation.

*PHABULOSA* (*PHB*), a member of the class III HD-ZIP gene family, leads cells toward adaxialization. Its dominant mutant, *phb-1d*, forms adaxialized radial leaves (31). Loss of function of *PHB* and other related class III HD-ZIP genes caused abaxialized radial cotyledons (32). Establishment of adaxial–abaxial polarity is required for leaf blade expansion, and loss of this polarity induces leaf radialization. The involvement of regulation of adaxial–abaxial polarity to heterophylly was reported in *Ranunculus trichophyllus* (6): submergence upregulated the genes *KANADIs*, which regulates abaxial growth, leading to the formation of abaxialized radial leaves. In the present study, expression of most of the adaxial–abaxial polarity genes, including *PHB*, was downregulated under submerged and ACC-treated conditions (Fig. 5F), suggesting that the loss of polarity is involved in the formation of submerged-type leaves.

### Blue light inhibits the submergence signal

In addition to the genes related to leaf morphogenesis, genes in the GO categories “response to light stimulus” and “shade avoidance” were affected by submergence (Fig. 4B). Submergence alters light quality via absorption and reflection in the water, and light quality influences various physiological responses in plants. We investigated how light quality influences leaf form under various light conditions. Blue light induced a pronounced response, causing the leaves to elongate along the anterior–posterior axis, and preventing leaflet narrowing in response to submergence (Fig. 6A). Based on our transcriptome analysis comparing the effects of white and blue light, the expression profile of submerged leaves under blue light was negatively correlated with that under white light (Fig. 6B), indicating that gene regulation normally induced during submergence under white light was not induced by submergence under blue light. In particular, the expression of submergence-induced ethylene response genes decreased under blue light (Fig. S6). Furthermore, the expression of abaxial–adaxial polarity regulating genes was not reduced under blue light. These findings indicate that the submergence signal was inhibited under blue light conditions via the ethylene response pathway.

**Figure 6.**
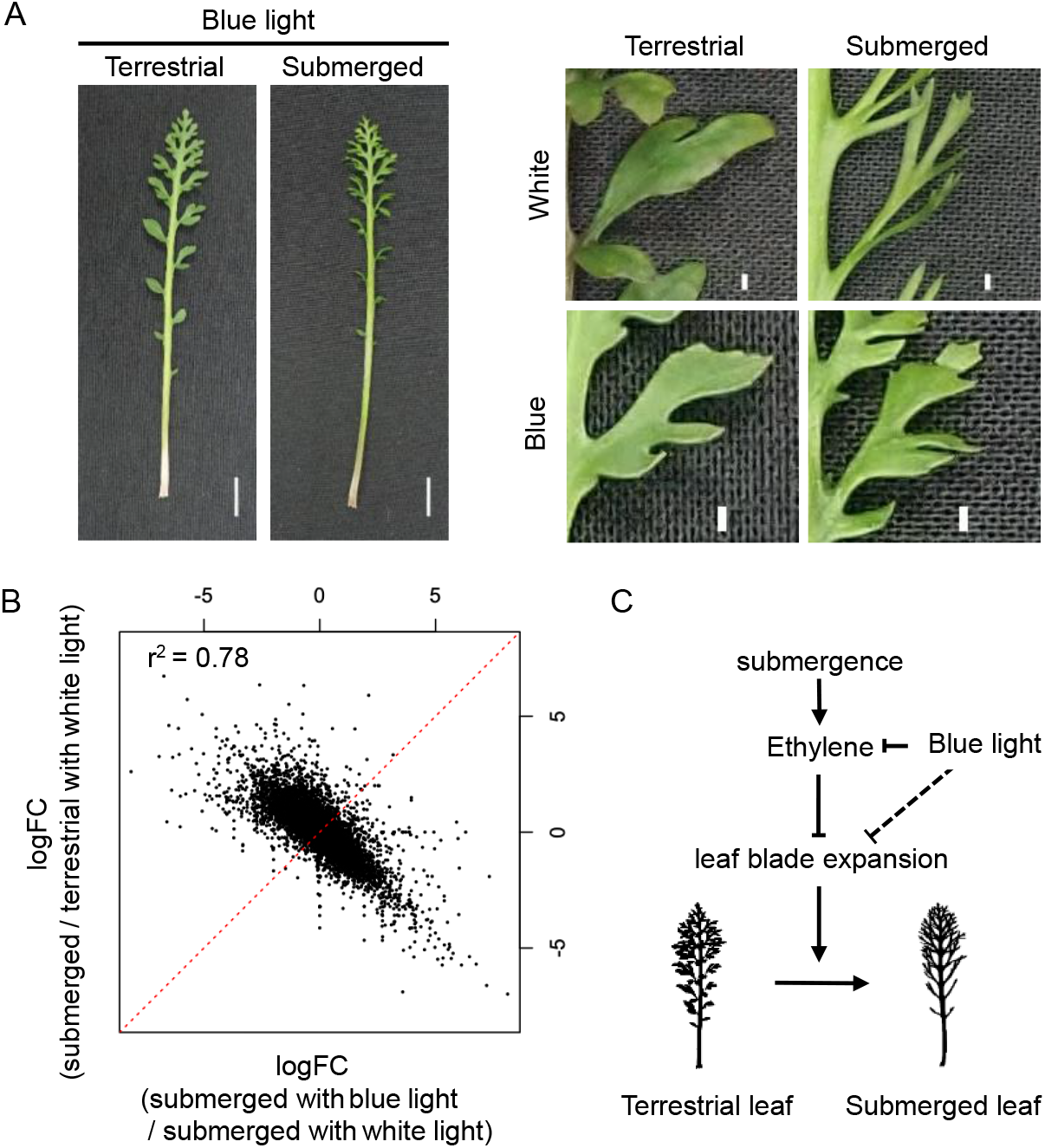
The effect of blue light on *Rorippa aquatica* heterophylly. (A) Mature leaves grown under blue light conditions (left panel; scale bar, 1 cm); magnified view of leaflet (right panel; scale bar, 1 mm). (B) Leaf transcriptome profile under white and blue light conditions. (C) Mechanistic model for heterophylly in response to submergence.

## Discussion

The chromosome-level genome assembly and comparative genomics analysis reveal the genome structure of *R. aquatica*, and elucidate its origin and evolution. We found that the *R. aquatica* genome originated by allotetraploidization through hybridization between two ancestral *Rorippa* species (Fig. 3). The hybridization occurred no earlier than 4.2 Mya, when *R. aquatica* subgenome A diverged from *R. islandica*. Thus, the genome of *R. aquatica* originated by hybridization between two *Rorippa* genomes with 8 chromosome pairs. The WGD was followed by post-polyploid descending dysploidy (from *n* = 16 to *n* = 15) mediated by nested chromosome insertion (NCI), and forming the fusion chromosome RaChr15. The NCI occurred independently in some population(s) of the tetraploid *Cardamine pratensis* (from 2*n* = 32 to 2*n* = 30) (33). Interestingly, both NCI events in *Cardamine* and *Rorippa* involved chromosome AK8/6 as a recipient chromosome, which recombined with chromosome AK2 in *R. aquatica* to form chromosome RaChr15, and chromosome AK5 in *C. pratensis*. Even more advanced post-polyploid descending dysploidy was documented in the tetraploid *C. cordifolia* where the chromosome number was reduced from 2*n* = 32 to 2*n* = 24 due to formation of five fusion chromosomes (34). On the contrary, the closely related tetraploid genomes (2*n* = 4x = 32) of horseradish (*Armoracia rusticana*) and watercress (*Nasturtium officinale*) contain structurally conserved parental subgenomes, except for a 2.4-Mb long unequal translocation in watercress (35). The *R. aquatica* genome sequence and assembly represent a genome-wide reference for future studies in *R. aquatica* and across the genus *Rorippa*.

Our transcriptome analysis of *R. aquatica*, based on whole genome assembly data, provides three key insights into heterophylly (Fig. 6C). First, the submergence signal was transmitted via ethylene, and the ethylene signaling inhibited leaf blade expansion. We found that ethylene signaling was induced by submergence, and exogenous ethylene resulted in narrower leaves even out of water. Since these responses have been reported in other amphibious plants (4, 7, 6) and ethylene signaling is a conserved pathway among angiosperms (36), it is potentially easily utilized as a submergence signal; this is consistent with its apparent role of regulating heterophylly in response to submergence in various plant species.

Gibberellins are also involved in regulation of *R. aquatica* leaf form, under both terrestrial and submerged conditions (3). Gibberellin treatment caused the emergence of simple leaves under low temperature and submergence, conditions normally inducing dissected leaves, and inhibiting the gibberellin signal caused dissected leaves even at high temperatures, inducing simple leaves. In the present study, however, the expression profile revealed increased gibberellin signaling in response to submergence. In other amphibious plants, which do not exhibit apparent heterophylly in response to temperature under the terrestrial condition, gibberellins exhibited different effects. Inhibiting the gibberellin signal suppresses the formation of the submerged-leaf phenotype, and gibberellin treatment fails to induce this leaf phenotype under terrestrial condition (4, 7). This suggests that *R. aquatica* leaf form might be regulated by two parallel pathways: temperature-dependent heterophylly mediated by gibberellins, and submergence-responsive heterophylly mediated by ethylene. How the two different pathways are regulated and interact is a subject for future work.

A second key insight of our study is that, in *R. aquatica*, both adaxial and abaxial genes were downregulated under submergence. This is similar to the situation in *Callitriche palustris* (7). The establishment of adaxial–abaxial polarity eventually leads to the establishment of the middle domain, which is situated at the juxtaposition between the adaxial–abaxial domains, and participates in leaf lamina outgrowth (37). Therefore, in *R. aquatica*, the suppression of leaf-blade expansion in submerged leaves may be due to changes in genes involved in establishing adaxial– abaxial polarity; such changes may prevent the establishment of the middle domain required for leaf blade outgrowth.

A third key insight of our study is that blue light is involved in regulating heterophylly. At the gene expression level, blue light blocked ethylene response-gene upregulation during submergence. Although the mechanism via which blue light regulates heterophylly through ethylene is unclear, the relationship between blue light and ethylene was studied in the shade avoidance response. For instance, in *A. thaliana*, low blue light induces stem elongation for shade avoidance, but not in ethylene-insensitive mutants (38). The molecular mechanisms underlying the blue light and ethylene response pathways are not clear, even in *A. thaliana*. In plants, blue-light reception is mediated by cryptochromes (CRYs) (39). In *Brassica napus*, overexpression of *CRY1* causes downregulation of the ethylene-biosynthesis-related genes *1-aminocyclopropane-1-carboxylate synthase* 5 and 8 (40). Under natural conditions, the quality of light reaching submerged plants changes with the water level, owing to differences in light-absorbance ratios. Changes in light quality might provide detailed signals about the underwater conditions. Our findings show that blue light plays an important role in regulating heterophylly in response to submergence. Considering that amphibious fern *Marsilea quadrifolia* showed similar response (41), blue light signaling may also be central to heterophylly in various plants. Elucidating blue light signaling may be key to elucidating heterophylly and its evolution.

The allotetraploid origin of *R. aquatica* suggests several possible mechanisms by which it attained traits such as amphibiousness and heterophylly in response to various signals. For instance, they could have arisen via inheritance from either parent or it could be a result of heterosis arising from crossing with other *Rorippa* species. Furthermore, they could have arisen from redundancy due to gene duplication, which often enables genes to acquire novel functions. Finally, the accelerated accumulation of mutations may have given rise to these traits. Future comparative studies of the species of origin of each subgenome is required to address these possibilities.

## Materials and methods

### Plant material

*Rorippa aquatica* plants (two accessions, N and S) (20) were kept in a growth chamber at 30 °C under continuous light at 50 µmol photons m^−2^ s^−1^ supplied by a fluorescent lamp. For each treatment, plants that regenerated from a leaf tip as described previously (42) were used.

To induce inflorescences for comparative chromosome painting, the growth chamber temperature for *R. aquatica* accession S was changed to 20 °C. Young inflorescences were collected from plants and fixed in freshly prepared fixative (ethanol: acetic acid, 3:1) overnight, transferred to 70% ethanol, and subsequently stored at −20 °C.

All plants used for morphological and transcriptome analyses were grown in a growth chamber at 25 °C. Plants which were used to examine the change of morphology and gene expression after transition to the submerged condition were grown in glass tanks with an approximate 8-cm water depth. To examine how ethylene affects heterophylly, all the plants were treated with 100 µM 1-aminocyclopropane-1-carboxylic acid (ACC) or 1 µM AgNO_3_. For chemical treatment under submerged conditions, the chemicals to be tested were diluted in the 200 mL sterile distilled water in the culture jar. All plants were grown under each treatment for two months, until the leaves were mature. To examine the effect of ethylene amount, plants were treated with different concentrations of ACC (10, 100 and 1000 µM) under the terrestrial condition as described above. To examine the effects of blue light, plants were grown under 20 µmol photons m^−2^ s^−1^ supplied by a blue LED.

### Chromosome preparation

Chromosome spreads from fixed young flower buds containing immature anthers were prepared according to published protocols (43, 44). Chromosome preparations were treated with 100 µg/mL RNase in 2× sodium saline citrate (SSC, 20× SSC: 3 M sodium chloride, 300 mM trisodium citrate, pH 7.0) for 60 min, and with 0.1 mg/mL pepsin in 0.01 M HCl at 37 °C for 5 min, then post-fixed in 4% formaldehyde in distilled water and dehydrated via an ethanol series (70%, 90%, and 100%, 2 min each).

### Painting probes

For comparative chromosome painting (CCP), 674 chromosome-specific BAC clones of *Arabidopsis thaliana* (The Arabidopsis Information Resource, TAIR; http://www.arabidopsis.org) were used to establish contigs corresponding to the 22 genomic blocks (GBs) and eight chromosomes of the Ancestral Crucifer Karyotype (ACK) (18). To determine and characterize inversions of GBs on chromosome Ra15, BAC contigs corresponding to GBs D and E were split into smaller subcontigs and differentially labelled to be used in several consecutive experiments. All DNA probes were labelled with biotin-dUTP, digoxigenin-dUTP, or Cy3-dUTP by nick translation, as per Mandáková & Lysak (45).

### Comparative chromosome painting

DNA probes were pooled appropriately, ethanol precipitated, dried, and dissolved in 20 μL of 50% formamide and 10% dextran sulfate in 2× SSC. The dissolved probe (20 μL) was pipetted onto a chromosome-containing slide and immediately denatured on a hot plate at 80 °C for 2 min. Hybridization was conducted in a moist chamber at 37 °C overnight. Post-hybridization washing was performed in 20% formamide in 2× SSC at 42 °C. Hybridized probes were visualized either as the direct fluorescence of Cy3-dUTP or via fluorescently labelled antibodies against biotin-dUTP and digoxigenin-dUTP (45). Chromosomes were counterstained with 4′,6-diamidino-2-phenylindole (DAPI, 2 µg/mL) in Vectashield antifade (Vector Laboratories). Fluorescence signals were analyzed and photographed using a Zeiss Axio Imager epifluorescence microscope with a CoolCube camera (MetaSystems, Altlussheim, Germany). Images were acquired separately for all four fluorochromes using appropriate excitation and emission filters (AHF Analysentechnik, Tübingen, Germany). The four monochromatic images were pseudocolored, merged, and cropped using Photoshop CS (Adobe Systems, Mountain View, CA) and ImageJ (National Institutes of Health, Bethesda, MA).

### Illumina genome DNA sequencing

Genome-seq libraries were constructed using whole- or nucleic-genome DNA. For extraction of nucleic DNA, the nuclear fraction was prepared from whole plants using the ‘Semi-pure Preparation of Nuclei Procedures’ protocol of the CelLytic PN Isolation/Extraction Kit (Sigma-Aldrich, St. Louis, MO). Genomic DNA was isolated from the nucleus or whole plant using a DNeasy Plant mini kit (Qiagen, Hilden, Germany). Genome-seq libraries were prepared using the Nextera DNA Sample Prep Kit. Sequencing was performed using NextSeq 500, generating paired-end reads of 151 bp.

### PacBio genome DNA sequencing

DNA for PacBio library was prepared as follows: crude nuclei were obtained from regenerated plants (using ca. 1 cm lengths of leaf tip) using the ‘Crude Preparation of Nuclei Procedures’ protocol of the CelLytic PN Isolation/Extraction Kit (Sigma-Aldrich). DNA extraction from crude nuclei was performed using two different methods. For the first run, the Dneasy Plant mini kit was used. For the subsequent two runs, genomic DNA was extracted using phenol/chloroform/isoamyl alcohol extraction with CTAB buffer and purified using QIAGEN Genomic-tip 20/G. Long reads were generated using the PacBio RS II system.

### Hi-C Seq

Preparation of the Hi-C Seq sample was performed as previously (46). HindIII was used for DNA digestion. For the preparation of the sequencing library, the purified Hi-C sample (500 ng) was diluted to 500 µl with dH_2_O, and 500 µl of 2× binding buffer (BB) (10 mM Tris, 1 mM EDTA, 2 M NaCl) was added. The diluted Hi-C samples were fragmented to a mean size of 300 bp by sonication using a Covaris M220 sonication system (Covaris, Woburn, MA, USA) in a milliTUBE 1 ml AFA Fibre (Covaris). The parameters of the program were as follows: power mode, frequency sweeping; time, 20 min; duty cycle, 5%; intensity, 4; cycles per burst, 200; temperature (water bath), 6 °C. Biotin-labelled Hi-C samples were then enriched using MyOne Streptavidin C1 magnetic beads (Veritas, Tokyo, Japan). For this, 60 µl of streptavidin beads were washed twice with 400 µl of Tween Wash Buffer (TWB) (5 mM Tris, 0.5 mM EDTA, 1M NaCl, 0.05% Tween-20). The recovery of streptavidin beads was performed by placing the tubes on a magnetic stand. Subsequently, the beads were added to 1 ml of sheared Hi-C sample. After 15 min of incubation at room temperature under rotation, the supernatant was removed, and the beads binding biotinylated Hi-C fragments were resuspended in 400 µl of 1× BB. Then, the beads were washed once in 60 µl RSB (Resuspension buffer) (Illumina, San Diego, CA, USA), and finally resuspended in 50 µl RSB. The enriched biotinylated DNA fragments were subjected to library construction on beads using the KAPA HyperPrep Kit for Illumina (Roche, Basel, Switzerland) according to the manufacturer’s protocol, with 18 cycles of PCR for library amplification. The amplified DNA fraction (50 µl) was corrected and purified using Agencourt AMPure XP (Beckman Coulter) following the standard protocol, and finally resuspended in 15 µl of RSB. The library was sequenced using a NextSeq 500 system, generating paired-end reads of 151 bp.

### Genome size estimation

The genome size of *R. aquatica* was estimated by k-mer counting using jellyfish2 (http://www.genome.umd.edu/jellyfish.html). K-mers from Illumina read data were counted, and the k-mer distribution was plotted; the distribution peaks from homozygous regions were picked manually, and genome size (in bases) was calculated as total number of k-mers / peak of k-mer distribution.

### Genome assembly and annotation

Genome assembly was performed using MaSuRCA (21) with both Illumina and PacBio reads. The assembled scaffolds were error-corrected using Pilon (47). Scaffolding into chromosome-level sequences was performed via the 3D de novo assembly (3D-DNA) pipeline (48), using the assembled scaffolds and Hi-C Seq reads. The remaining gaps in chromosome-level sequences were filled by LR_Gapcloser (49), using PacBio reads that were error-corrected using ColorMap (50). Assembled genome sequences were benchmarked using Benchmarking Universal Single-Copy Orthologs (BUSCO) (51) with a land-plant dataset (embryophyta_odb9). Repeat sequences in the genome were identified and masked using RepeatModeler and RepeatMasker (http://www.repeatmasker.org). Gene prediction was performed using the PASA pipeline (23). Three types of prediction were used: 1) ab initio prediction using AUGUSTUS (52), GlimmerHMM (53), and SNAP, with an *Arabidopsis* training dataset; 2) Protein homology detection using EXONERATE with *A. thaliana* TAIR10 protein data; and 3) Alignment of assembled transcripts to the genome. Transcriptome data were obtained by de novo assembly using Trinity (54), with RNAseq data (DRA006777) from a published paper (42). All the predicted gene structures were integrated into the final gene data using EvidenceModeler (EVM) (55) and PASA. Gene Ontology terms were assigned to each transcript using Blast2GO (56) based on the results of a BLASTP homology search against the non-redundant protein sequence (Nr) database and InterProScan.

### Genome structure

Alignment of *R. aquatica* chromosome sequences to the *C. hirsuta* genome was performed using MUMMER (57). Genome structure (distribution of genes and long terminal repeats, and links between paralogous genes) was illustrated using CIRCOS (58).

### Comparative genomics analysis

We performed whole-genome level phylogenetic analysis using *R. aquatica* genomic information and genome-level data of several plant species. The protein dataset of 22 plant species from the Phytozome database (https://phytozome-next.jgi.doe.gov/). The datasets of *C. hirsuta* and *Barbarea vulgaris* were prepared using data from each species’ genome database (http://chi.mpipz.mpg.de/ (59) and http://plen.ku.dk/Barbarea (60), respectively). We prepared draft *Rorippa islandica* genomic data via genome assembly using Velvet (61) with genome-seq read data (SRR1801303) from the Sequence Read Archive, setting k-mer to 151. Gene prediction was performed using AUGUSTUS (52) using an *Arabidopsis* training dataset. Protein sequences of a single representative longest transcript variant for each gene were extracted using an inhouse Perl script. Using OrthoFinder (62), each protein sequence was clustered into an orthogroup based on similarity, and the phylogenetic analysis of each orthogroup was integrated as a species tree.

The synonymous substitution rate (Ks) was calculated to estimate evolutionary event ages. Using MACSE (63), we performed multiple-alignment of the coding DNA sequences (CDS) in each orthogroups in which single-copy conserved genes in seven related Brassicaceae species (*Eutrema salsugineum, Arabidopsis thaliana, Arabidopsis lyrata, Barbarea vulgaris, Cardamine hirsuta, Capsella rubella*, and *Rorippa islandica*) and duplicated genes in *R. aquatica* were classified, then we calculated Ks using yn00 in the PAML package (http://abacus.gene.ucl.ac.uk/software/paml.html). The age of each event was estimated as T (in years) = Ks of peak / (2 * *μ*), where *μ*, the synonymous divergence rate per site per year, equals 6.51648E−09 in Brassicaceae (26).

### Transcriptome analysis

Total RNA was isolated from shoot apexes containing young leaves using RNeasy Plant Mini kit (QIAGEN, Hilden, Germany). RNAseq libraries were prepared using the Illumina TruSeq Stranded RNA LT kit (Illumina, CA, USA), according to the manufacturer’s instructions. Libraries were sequenced on the NextSeq500 sequencing platform (Illumina, CA, USA), and 76 bp single-end reads were obtained. The reads were mapped to the genome sequences of *R. aquatica* using Tophat2. Count data were subjected to a trimmed mean of M-value normalization in edgeR (64). Transcript expression and DEGs were defined using the edgeR GLM approach.

## Supporting information

Supplementary Tabels

## Data Availability

The assembled *R. aquatica* genome sequences and its annotations have been deposited in Figshare (10.6084/m9.figshare.19207362). Genome-seq read data and Hi-C seq read data are available in the DDBJ Sequenced Read Archive (DRA) under the accession numbers DRA010675 and DRA013596, respectively. Transcriptome read data are also available in DDBJ DRA under DRA014113, DRA014114, DRA014164, and DRA014165.

## Acknowledgements

We thank Dr. Neelima Sinha, Dr. Naomi Nakayama, and Dr. Dhanya Radhakrishnan for useful discussions. This work was financially supported by the JSPS KAKENHI grants 21H02513 and MEXT-Supported Program for the Strategic Research Foundation at Private Universities grant S1511023 to S. K. and JSPS KAKENHI grants 20H05911 and 22H00415 to S. M. This work was also supported by the National Key Research and Development Program of China (2017YFE0128800), the National Natural Science Foundation of China (31870384, 32101254), and the International Partnership Program of the Chinese Academy of Sciences (152342KYSB20200021) to H. H. and G. L.

## References

1. H. Nakayama, N. R. Sinha, S. Kimura, How Do Plants and Phytohormones Accomplish Heterophylly, Leaf Phenotypic Plasticity, in Response to Environmental Cues. Front. Plant Sci. 8, 1717 (2017).

2. G. Li, S. Hu, H. Hou, S. Kimura, Heterophylly: Phenotypic Plasticity of Leaf Shape in Aquatic and Amphibious Plants. Plants Basel Switz. 8, E420 (2019).

3. H. Nakayama, et al., Regulation of the KNOX-GA gene module induces heterophyllic alteration in North American lake cress. Plant Cell 26, 4733–4748 (2014).

4. G. Li, et al., Water-Wisteria as an ideal plant to study heterophylly in higher aquatic plants. Plant Cell Rep. 36, 1225–1236 (2017).

5. G. Li, et al., Establishment of an Agrobacterium mediated transformation protocol for the detection of cytokinin in the heterophyllous plant Hygrophila difformis (Acanthaceae). Plant Cell Rep. 39, 737–750 (2020).

6. J. Kim, et al., A molecular basis behind heterophylly in an amphibious plant, Ranunculus trichophyllus. PLoS Genet. 14, e1007208 (2018).

7. H. Koga, M. Kojima, Y. Takebayashi, H. Sakakibara, H. Tsukaya, Identification of the unique molecular framework of heterophylly in the amphibious plant Callitriche palustris L. Plant Cell 33, 3272–3292 (2021).

8. C. La Rue, Regeneration in Radicula aquatica. Pap. Mich. Acad. Sci. Arts Lett. 28, 51–61 (1943).

9. G. Li, et al., Mechanisms of the Morphological Plasticity Induced by Phytohormones and the Environment in Plants. Int. J. Mol. Sci. 22, E765 (2021).

10. D. H. Les, Molecular systematics and taxonomy of lake cress (Neobeckia aquatica; Brassicaceae), an imperiled aquatic mustard. Aquat. Bot. 49, 149–165 (1994).

11. A. S. Hay, et al., Cardamine hirsuta: a versatile genetic system for comparative studies. Plant J. Cell Mol. Biol. 78, 1–15 (2014).

12. M. Bar, N. Ori, Compound leaf development in model plant species. Curr. Opin. Plant Biol. 23, 61–69 (2015).

13. H. Nakayama, K. Fukushima, T. Fukuda, J. Yokoyama, S. Kimura, Molecular Phylogeny Determined Using Chloroplast DNA Inferred a New Phylogenetic Relationship of Rorippa aquatica (Eaton) EJ Palmer & Steyermark (Brassicaceae)—Lake Cress. Am. J. Plant Sci. 05, 48–54 (2014).

14. S. I. Warwick, A. Francis, I. A. Al-Shehbaz, Brassicaceae: Species checklist and database on CD-Rom. Plant Syst. Evol. 259, 249–258 (2006).

15. J. G. King, J. D. Hadfield, The evolution of phenotypic plasticity when environments fluctuate in time and space. Evol. Lett. 3, 15–27 (2019).

16. E. Lafuente, P. Beldade, Genomics of Developmental Plasticity in Animals. Front. Genet. 10, 720 (2019).

17. M. E. Schranz, M. A. Lysak, T. Mitchell-Olds, The ABC’s of comparative genomics in the Brassicaceae: building blocks of crucifer genomes. Trends Plant Sci. 11, 535–542 (2006).

18. M. A. Lysak, T. Mandáková, M. E. Schranz, Comparative paleogenomics of crucifers: ancestral genomic blocks revisited. Curr. Opin. Plant Biol. 30, 108–115 (2016).

19. T. Mandáková, et al., The story of promiscuous crucifers: origin and genome evolution of an invasive species, Cardamine occulta (Brassicaceae), and its relatives. Ann. Bot. 124, 209–220 (2019).

20. H. Nakayama, et al., Comparative transcriptomics with self-organizing map reveals cryptic photosynthetic differences between two accessions of North American Lake cress. Sci. Rep. 8, 3302 (2018).

21. A. V. Zimin, et al., Hybrid assembly of the large and highly repetitive genome of Aegilops tauschii, a progenitor of bread wheat, with the MaSuRCA mega-reads algorithm. Genome Res. 27, 787–792 (2017).

22. B. J. Haas, et al., Improving the Arabidopsis genome annotation using maximal transcript alignment assemblies. Nucleic Acids Res. 31, 5654–5666 (2003).

23. B. J. Haas, et al., Automated eukaryotic gene structure annotation using EVidenceModeler and the Program to Assemble Spliced Alignments. Genome Biol. 9, R7 (2008).

24. S. M. Jeelani, S. Rani, S. Kumar, S. Kumari, R. C. Gupta, Cytological studies of Brassicaceae burn. (Cruciferae juss.) from Western Himalayas. Tsitol. Genet. 47, 26–36 (2013).

25. C.-H. Huang, et al., Resolution of Brassicaceae Phylogeny Using Nuclear Genes Uncovers Nested Radiations and Supports Convergent Morphological Evolution. Mol. Biol. Evol. 33, 394–412 (2016).

26. A. R. De La Torre, Z. Li, Y. Van de Peer, P. K. Ingvarsson, Contrasting Rates of Molecular Evolution and Patterns of Selection among Gymnosperms and Flowering Plants. Mol. Biol. Evol. 34, 1363–1377 (2017).

27. D. Wanke, The ABA-mediated switch between submersed and emersed life-styles in aquatic macrophytes. J. Plant Res. 124, 467–475 (2011).

28. A. Kuwabara, K. Ikegami, T. Koshiba, T. Nagata, Effects of ethylene and abscisic acid upon heterophylly in Ludwigia arcuata (Onagraceae). Planta 217, 880–887 (2003).

29. Y. K. Lee, et al., LONGIFOLIA1 and LONGIFOLIA2, two homologous genes, regulate longitudinal cell elongation in Arabidopsis. Dev. Camb. Engl. 133, 4305–4314 (2006).

30. D. Vlad, et al., Leaf shape evolution through duplication, regulatory diversification, and loss of a homeobox gene. Science 343, 780–783 (2014).

31. Y. Eshed, A. Izhaki, S. F. Baum, S. K. Floyd, J. L. Bowman, Asymmetric leaf development and blade expansion in Arabidopsis are mediated by KANADI and YABBY activities. Dev. Camb. Engl. 131, 2997–3006 (2004).

32. J. F. Emery, et al., Radial patterning of Arabidopsis shoots by class III HD-ZIP and KANADI genes. Curr. Biol. CB 13, 1768–1774 (2003).

33. T. Mandáková, et al., The more the merrier: recent hybridization and polyploidy in Cardamine. Plant Cell 25, 3280–3295 (2013).

34. T. Mandáková, A. D. Gloss, N. K. Whiteman, M. A. Lysak, How diploidization turned a tetraploid into a pseudotriploid. Am. J. Bot. 103, 1187–1196 (2016).

35. T. Mandáková, M. A. Lysak, Healthy Roots and Leaves: Comparative Genome Structure of Horseradish and Watercress. Plant Physiol. 179, 66–73 (2019).

36. B. M. Binder, Ethylene signaling in plants. J. Biol. Chem. 295, 7710–7725 (2020).

37. M. Nakata, et al., Roles of the middle domain-specific WUSCHEL-RELATED HOMEOBOX genes in early development of leaves in Arabidopsis. Plant Cell 24, 519–535 (2012).

38. R. Pierik, G. C. Whitelam, L. A. C. J. Voesenek, H. de Kroon, E. J. W. Visser, Canopy studies on ethylene-insensitive tobacco identify ethylene as a novel element in blue light and plant-plant signalling. Plant J. Cell Mol. Biol. 38, 310–319 (2004).

39. C. Lin, Plant blue-light receptors. Trends Plant Sci. 5, 337–342 (2000).

40. P. Sharma, M. Chatterjee, N. Burman, J. P. Khurana, Cryptochrome 1 regulates growth and development in Brassica through alteration in the expression of genes involved in light, phytohormone and stress signalling. Plant Cell Environ. 37, 961–977 (2014).

41. B. L. Lin, W. J. Yang, Blue light and abscisic acid independently induce heterophyllous switch in Marsilea quadrifolia. Plant Physiol. 119, 429–434 (1999).

42. R. Amano, et al., Molecular Basis for Natural Vegetative Propagation via Regeneration in North American Lake Cress, Rorippa aquatica (Brassicaceae). Plant Cell Physiol. 61, 353– 369 (2020).

43. M. A. Lysak, T. Mandáková, Analysis of plant meiotic chromosomes by chromosome painting. Methods Mol. Biol. Clifton NJ 990, 13–24 (2013).

44. T. Mandáková, M. A. Lysak, Chromosome Preparation for Cytogenetic Analyses in Arabidopsis. Curr. Protoc. Plant Biol. 1, 43–51 (2016).

45. T. Mandáková, M. A. Lysak, Painting of Arabidopsis Chromosomes with Chromosome-Specific BAC Clones. Curr. Protoc. Plant Biol. 1, 359–371 (2016).

46. S. Grob, U. Grossniklaus, Chromatin Conformation Capture-Based Analysis of Nuclear Architecture. Methods Mol. Biol. Clifton NJ 1456, 15–32 (2017).

47. B. J. Walker, et al., Pilon: An Integrated Tool for Comprehensive Microbial Variant Detection and Genome Assembly Improvement. PLOS ONE 9, e112963 (2014).

48. O. Dudchenko, et al., De novo assembly of the Aedes aegypti genome using Hi-C yields chromosome-length scaffolds. Science 356, 92–95 (2017).

49. G.-C. Xu, et al., LR_Gapcloser: a tiling path-based gap closer that uses long reads to complete genome assembly. GigaScience 8 (2019).

50. E. Haghshenas, F. Hach, S. C. Sahinalp, C. Chauve, CoLoRMap: Correcting Long Reads by Mapping short reads. Bioinforma. Oxf. Engl. 32, i545–i551 (2016).

51. F. A. Simão, R. M. Waterhouse, P. Ioannidis, E. V. Kriventseva, E. M. Zdobnov, BUSCO: assessing genome assembly and annotation completeness with single-copy orthologs. Bioinforma. Oxf. Engl. 31, 3210–3212 (2015).

52. M. Stanke, S. Waack, Gene prediction with a hidden Markov model and a new intron submodel. Bioinforma. Oxf. Engl. 19 Suppl 2, ii215–225 (2003).

53. S. L. Salzberg, M. Pertea, A. L. Delcher, M. J. Gardner, H. Tettelin, Interpolated Markov models for eukaryotic gene finding. Genomics 59, 24–31 (1999).

54. M. G. Grabherr, et al., Full-length transcriptome assembly from RNA-Seq data without a reference genome. Nat. Biotechnol. 29, 644–652 (2011).

55. J. E. Allen, M. Pertea, S. L. Salzberg, Computational gene prediction using multiple sources of evidence. Genome Res. 14, 142–148 (2004).

56. A. Conesa, S. Götz, Blast2GO: A comprehensive suite for functional analysis in plant genomics. Int. J. Plant Genomics 2008, 619832 (2008).

57. S. Kurtz, et al., Versatile and open software for comparing large genomes. Genome Biol. 5, R12 (2004).

58. M. I. Krzywinski, et al., Circos: An information aesthetic for comparative genomics. Genome Res. (2009) https://doi.org/10.1101/gr.092759.109 (December 6, 2021).

59. X. Gan, et al., The Cardamine hirsuta genome offers insight into the evolution of morphological diversity. Nat. Plants 2, 16167 (2016).

60. S. L. Byrne, et al., The genome sequence of Barbarea vulgaris facilitates the study of ecological biochemistry. Sci. Rep. 7, 40728 (2017).

61. D. R. Zerbino, E. Birney, Velvet: algorithms for de novo short read assembly using de Bruijn graphs. Genome Res. 18, 821–829 (2008).

62. D. M. Emms, S. Kelly, OrthoFinder: solving fundamental biases in whole genome comparisons dramatically improves orthogroup inference accuracy. Genome Biol. 16, 157 (2015).

63. V. Ranwez, S. Harispe, F. Delsuc, E. J. P. Douzery, MACSE: Multiple Alignment of Coding SEquences Accounting for Frameshifts and Stop Codons. PLOS ONE 6, e22594 (2011).

64. M. D. Robinson, D. J. McCarthy, G. K. Smyth, edgeR: a Bioconductor package for differential expression analysis of digital gene expression data. Bioinforma. Oxf. Engl. 26, 139–140 (2010).

