## Supplementary Figures for "Chromosome-level genome assembly of *Rorippa aquatica* revealed its allotetraploid origin and mechanisms of heterophylly upon submergence"

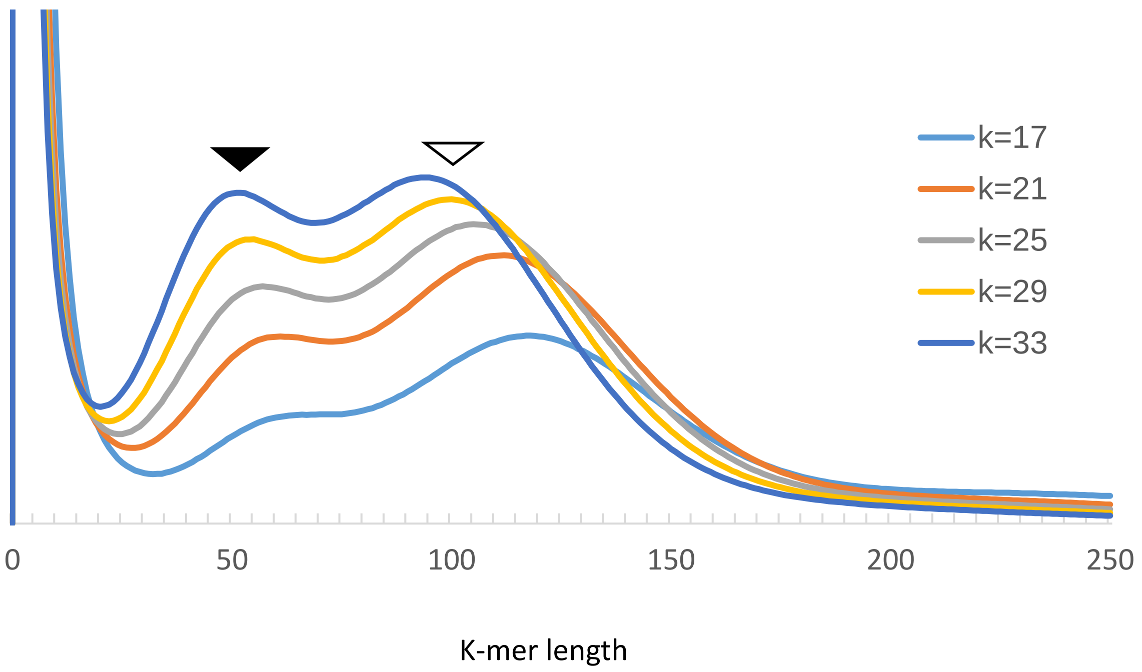


Figure S1. K-mer distribution of genome-seq reads of *R. aquatica*. Solid and open arrowhead indicates K-mer peak from heterozygous and homozygous region, respectively.

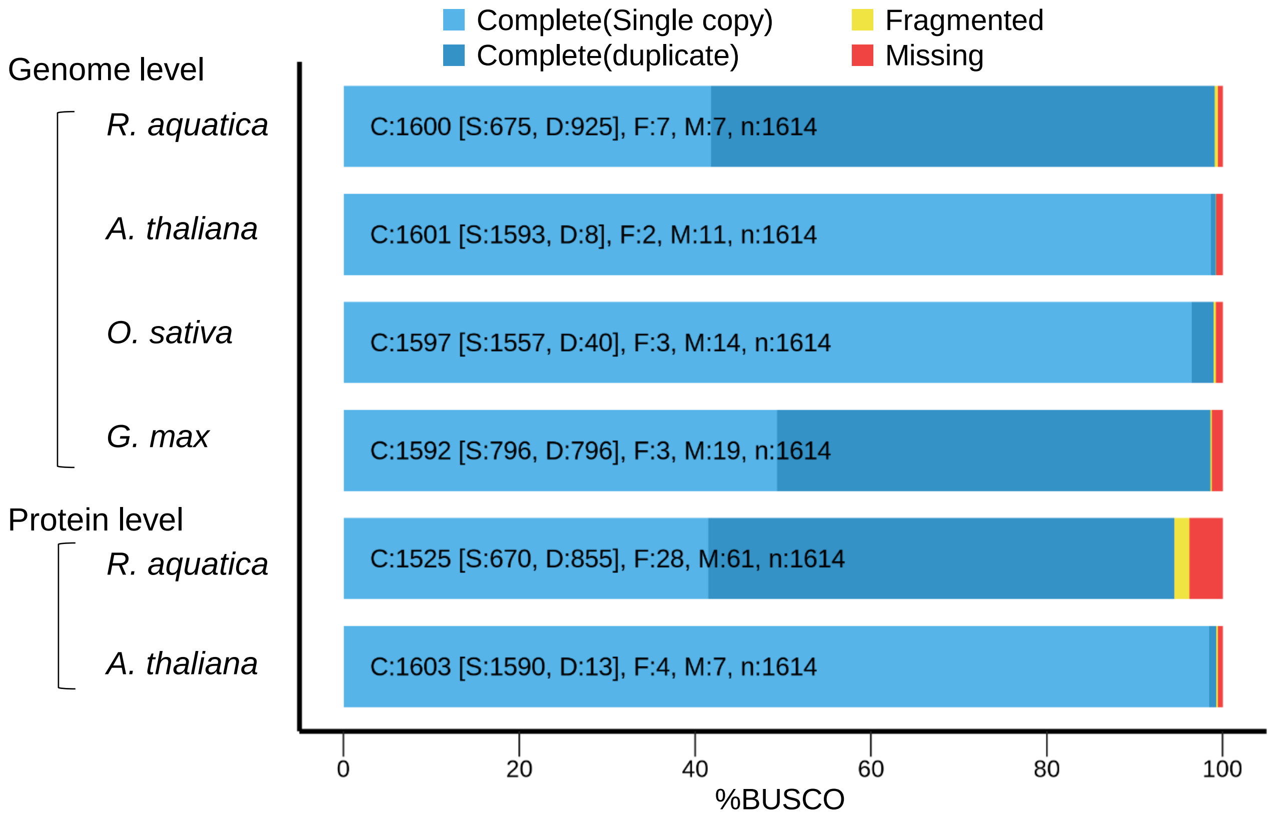


Figure S2. Quality assessment of *R. aquatica* genome by BUSCO.

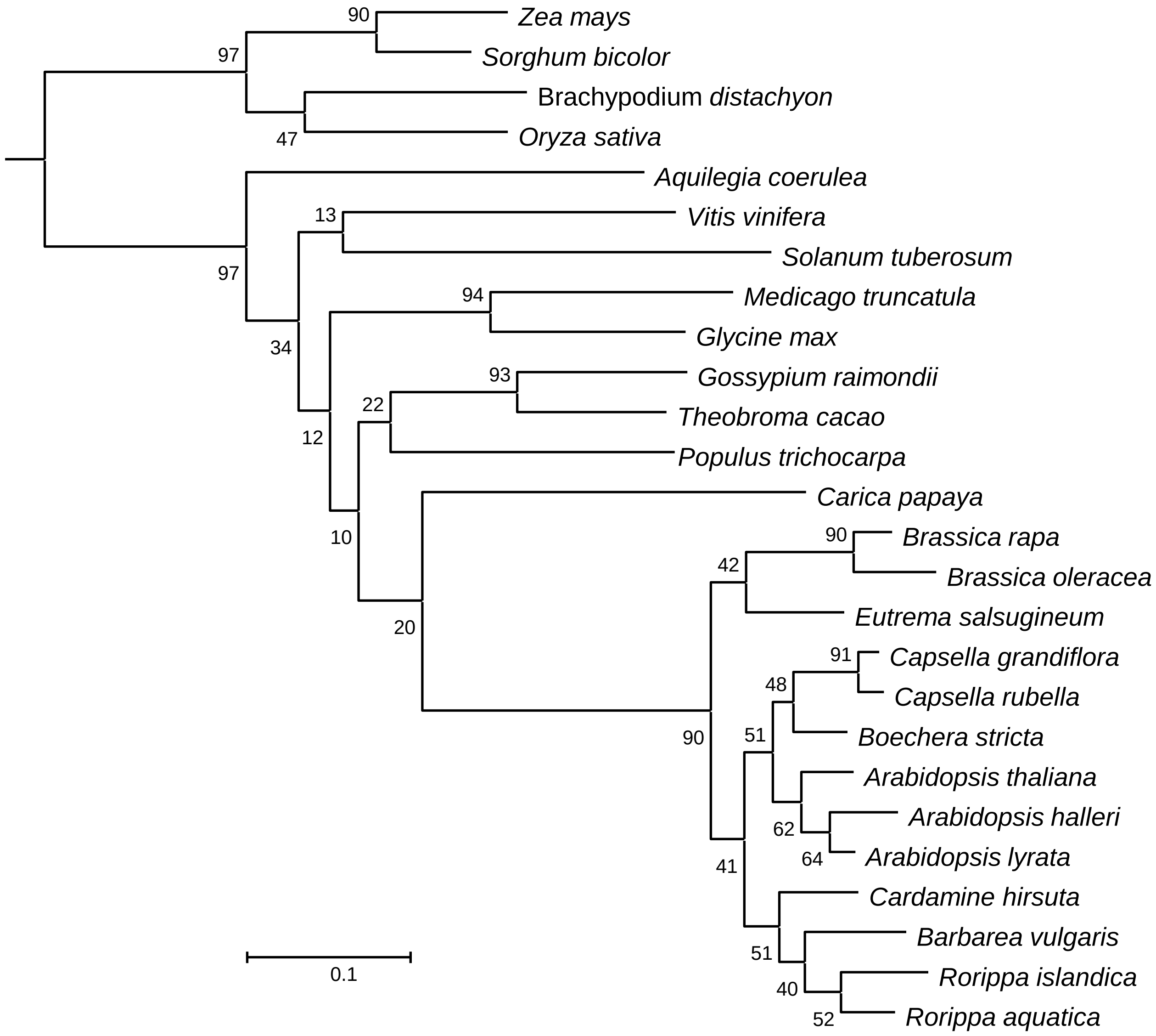


Figure S3. Phylogenetic tree from whole-genome level data. Scale bar indicates branch length (substitutions per site).

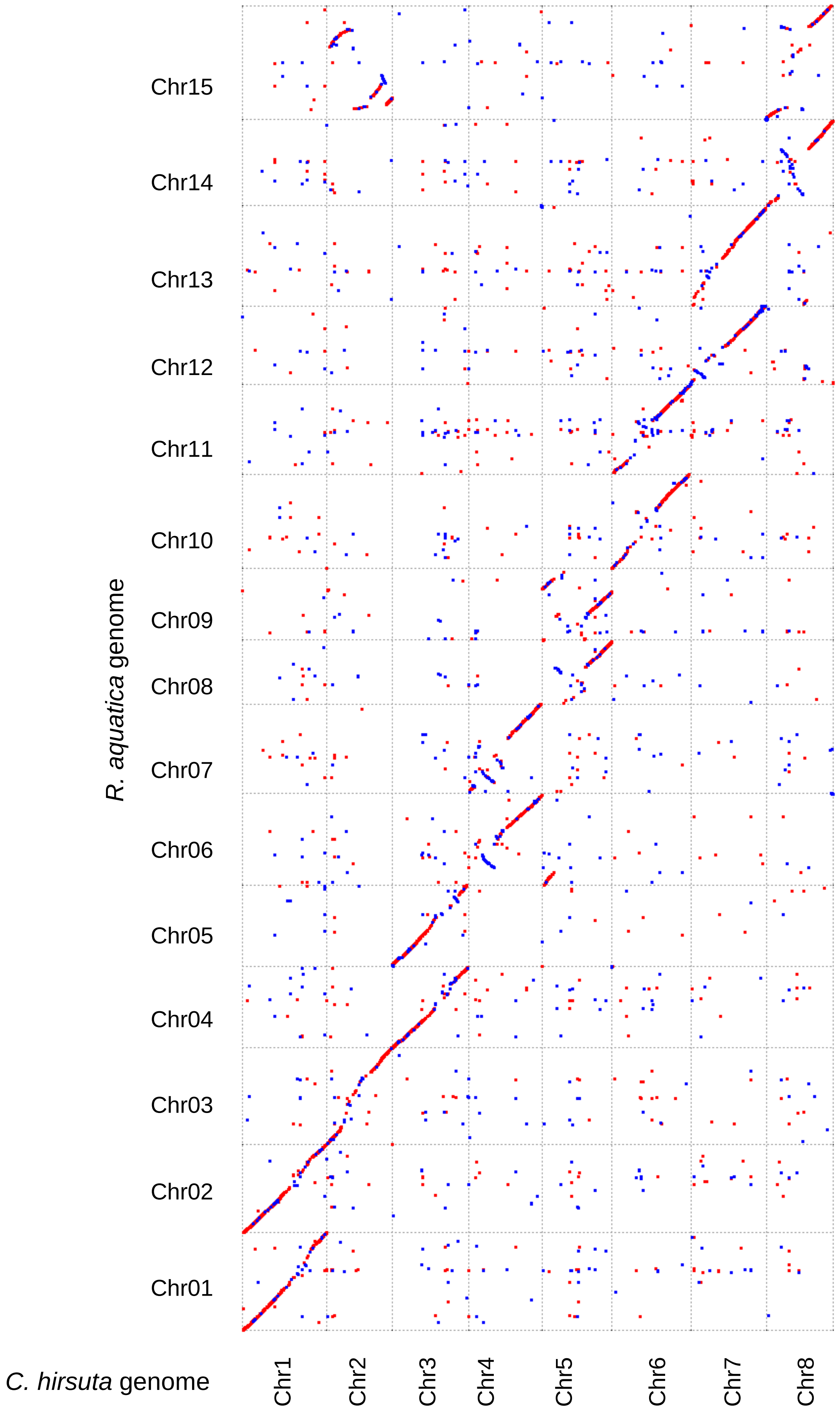


Figure S4. Comparison of genome sequences between *Cardamine hirsuta* and *Rorippa aquatica*. Dots with red and blue indicate the region that show similarity to forward and reverse strand of *C. hirsuta* genome respectively.

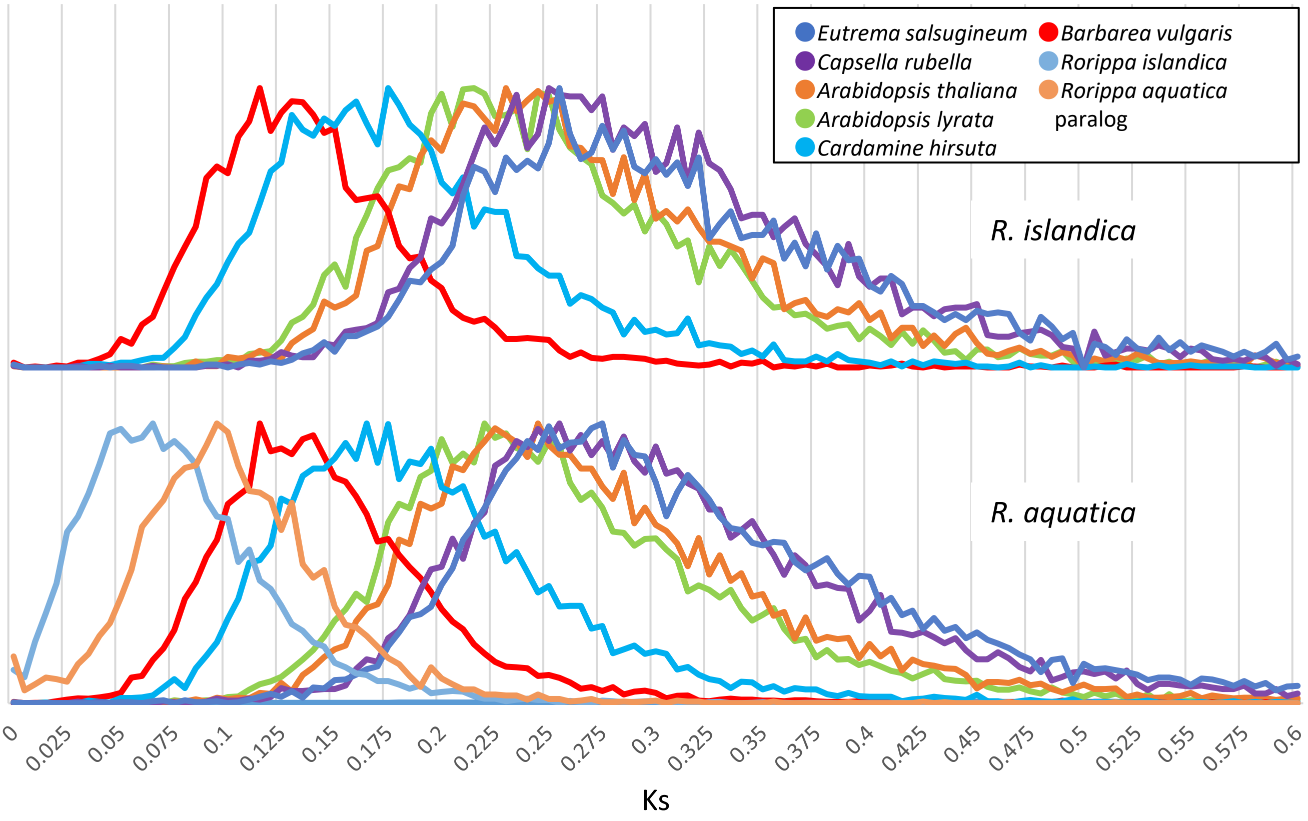


Figure S5. Ks distribution among genes that are conserved as single copy in most Brassicaceae species but duplicated in *R. aquatica.*

A

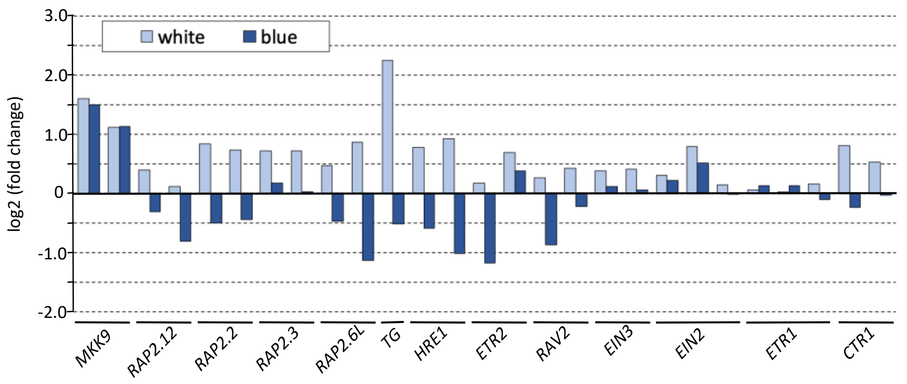


B

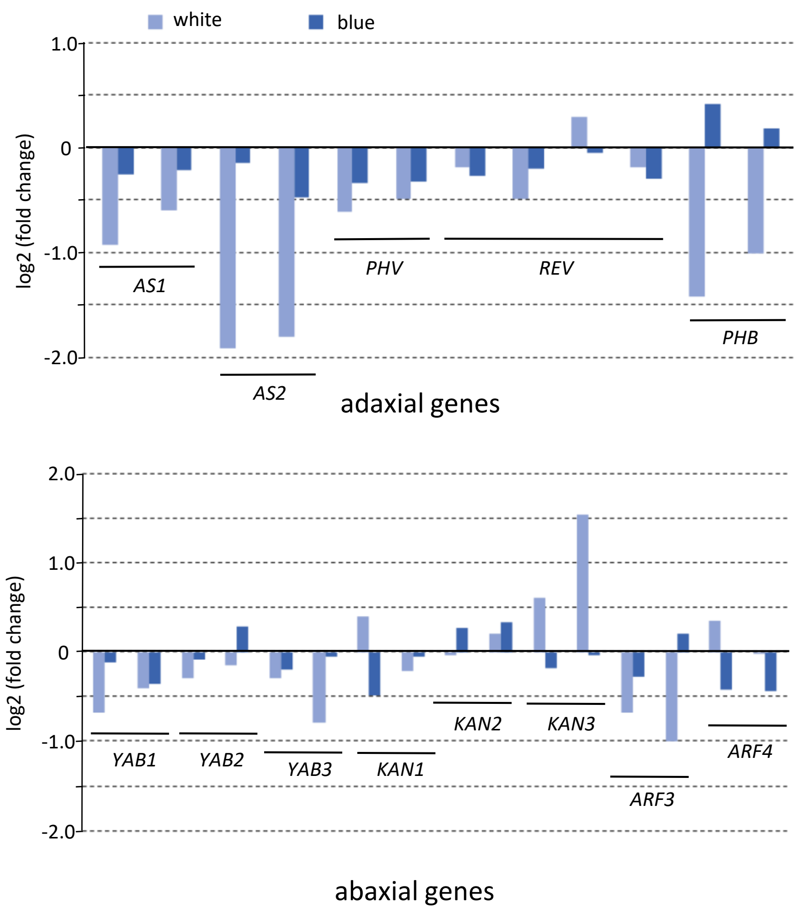


Figure S6. Expression profile of ethylene response genes (A) and adaxial-abaxial polarity genes (B) after transfer to submergence under white or blue light condition. Graphs depict fold change at 1 hour after transfer from terrestrial condition under white light to submerged condition under white or blue light.
