## Supplementary Tabels for "Chromosome-level genome assembly of *Rorippa aquatica* revealed its allotetraploid origin and mechanisms of heterophylly upon submergence"

Table S1. Summary of assembled R. *aquatica* genome sequences.

| | |
|---|---|
| Number of Chromosomal sequences | 15 |
| Total number of sequences | 2,055 |
| Num. of sequences (>= 50kbp) | 120 |
| Total length（bp） | 452,230,509 |
| Total length (>= 50kbp) | 414,395,297 |
| Total length (Chromosomes) | 414,395,297 |
| Length of longest sequence | 35,558,744 |
| GC (%) | 35.31 |
| N50 | 28,163,506 |

Table S2. Summary of genome size estimation by k-mer counting.

| | k=17 | k=21 | k=25 | k=29 | k=33 |
|---|---|---|---|---|---|
| total number of k-mers ($x10^6$) | 49,594 | 47,728 | 45,964 | 44,055 | 42,019 |
| peak of k-mer distribution | 117 | 111 | 105 | 100 | 95 |
| estimated genome size (Mb) | 423.89 | 429.99 | 437.76 | 440.55 | 442.31 |

Table S3. Statistics of repeat masking.

| | Number of element | Length | % of genome |
|---|---|---|---|
| SINEs | 1,801 | 354,676 | 0.08 |
| LINEs | 9,692 | 9,767,096 | 2.16 |
| LTR_elements | 120,041 | 133,198,170 | 29.45 |
| DNA_elements | 57,852 | 22,871,963 | 5.06 |
| Unclassified | 85,028 | 42,031,712 | 9.29 |
| total | 274,414 | 208,223,617 | 46.04 |

Table S4. Statistics of predicted gene structure.

| | transcript | gene |
|---|---|---|
| number | 94666 | 46197 |
| max length | 16567 | 16567 |
| Minimum length | 150 | 150 |
| average length | 1912.4 | 1458.7 |
| median length | 1775 | 1266 |

Table S5. Gene distribution in 2 *Rorippa* species classified into Brassicaceae conserved single gene orthogroup.

| Gene number in orthogroup | *R. aquatica* number | *R. aquatica* ratio(%) | *R. islandica* number | *R. islandica* ratio(%) |
|---|---|---|---|---|
| 0 | 529 | 4.9 | 449 | 4.1 |
| 1 | 3682 | 34.0 | 9896 | 91.2 |
| 2 | 6355 | 58.6 | 407 | 3.8 |
| 3 | 252 | 2.3 | 71 | 0.7 |
| 4 | 21 | 0.2 | 17 | 0.2 |
| >=5 | 6 | 0.1 | 5 | 0.0 |
| Total | 10845 | 100.0 | 10845 | 100.0 |

Table S6. Divergence time estimated from Ks. Age estimates given as million years ago.

| | *R. aquatica* | | | | *R. islandica* | | | |
|---|---|---|---|---|---|---|---|---|
| | Ks | | age | | Ks | | age | |
| | average | median | average | median | average | median | average | median |
| *E. salsugineum* | 0.331 | 0.304 | 25.4 | 23.3 | 0.326 | 0.299 | 25.0 | 22.9 |
| *C. rubella* | 0.316 | 0.296 | 24.3 | 22.7 | 0.311 | 0.291 | 23.8 | 22.3 |
| *A. thaliana* | 0.281 | 0.264 | 21.5 | 20.2 | 0.275 | 0.257 | 21.1 | 19.7 |
| *A. lyrata* | 0.266 | 0.250 | 20.4 | 19.2 | 0.260 | 0.245 | 20.0 | 18.8 |
| *C. hirsuta* | 0.196 | 0.185 | 15.0 | 14.2 | 0.189 | 0.178 | 14.5 | 13.7 |
| *B. vulgaris* | 0.151 | 0.142 | 11.6 | 10.9 | 0.146 | 0.137 | 11.2 | 10.5 |
| *R. islandica* | 0.081 | 0.073 | 6.2 | 5.6 | | | | |
| *R. aquatica* paralogs | 0.108 | 0.102 | 8.3 | 7.8 | | | | |

Table S7. Median Ks values at chromosome level. Letters at median show groups assigned based on fold discovery rate (FDR) by pairwise wilcoxon rank sum tests (FDR < 0.05)

| Subgenome A group | | Subgenome B group | |
|---|---|---|---|
| RaChr01 | 0.0526 $^{ab}$ | RaChr02 | 0.0884 $^{d}$ |
| RaChr05 | 0.0564 $^{bc}$ | RaChr04 | 0.0954 $^{ef}$ |
| RaChr07 | 0.0551 $^{bc}$ | RaChr06 | 0.0953 $^{ef}$ |
| RaChr08 | 0.0522 $^{a}$ | RaChr09 | 0.0920 $^{de}$ |
| RaChr10 | 0.0579 $^{c}$ | RaChr11 | 0.0931 $^{de}$ |
| RaChr13 | 0.0529 $^{ab}$ | RaChr12 | 0.0908 $^{de}$ |
| RaChr03 | 0.0529 $^{ab}$ | RaChr15(03ortho) | 0.0987 $^{f}$ |
| RaChr14 | 0.0535 $^{ab}$ | RaChr15(14ortho) | 0.0946 $^{ef}$ |
| Total | 0.0541 | Total | 0.0935 |

Table S8. Statistics analysis of Ks by pairwise wilcoxon rank sum tests. Values show FDR. Pale blue and orange indicates significant difference with FDR < 0.05 and <2e-16, respectively.

| | Subgenome A | | | | | | | | Subgenome B | | | | | | |
|---|---|---|---|---|---|---|---|---|---|---|---|---|---|---|---|
| | RaChr01 | RaChr03 | RaChr05 | RaChr07 | RaChr08 | RaChr10 | RaChr13 | RaChr14 | RaChr02 | RaChr04 | RaChr06 | RaChr09 | RaChr11 | RaChr12 | RaChr15 (03ortho) |
| **RaChr01** | - | - | - | - | - | - | - | - | - | - | - | - | - | - | - |
| **RaChr03** | 0.80385 | - | - | - | - | - | - | - | - | - | - | - | - | - | - |
| **RaChr05** | 0.16474 | 0.37411 | - | - | - | - | - | - | - | - | - | - | - | - | - |
| **RaChr07** | 0.10002 | 0.28505 | 0.72836 | - | - | - | - | - | - | - | - | - | - | - | - |
| **RaChr08** | 0.29485 | 0.26331 | 0.02451 | 0.01562 | - | - | - | - | - | - | - | - | - | - | - |
| **RaChr10** | 0.00155 | 0.02895 | 0.1434 | 0.34319 | 0.00029 | - | - | - | - | - | - | - | - | - | - |
| **RaChr13** | 0.89277 | 0.72836 | 0.14954 | 0.10049 | 0.39479 | 0.00253 | - | - | - | - | - | - | - | - | - |
| **RaChr14** | 0.92087 | 0.80385 | 0.30465 | 0.18251 | 0.3077 | 0.01454 | 0.83646 | - | - | - | - | - | - | - | - |
| **RaChr02** | < 2e-16 | < 2e-16 | < 2e-16 | < 2e-16 | < 2e-16 | < 2e-16 | < 2e-16 | < 2e-16 | - | - | - | - | - | - | - |
| **RaChr04** | < 2e-16 | < 2e-16 | < 2e-16 | < 2e-16 | < 2e-16 | < 2e-16 | < 2e-16 | < 2e-16 | 0.00826 | - | - | - | - | - | - |
| **RaChr06** | < 2e-16 | < 2e-16 | < 2e-16 | < 2e-16 | < 2e-16 | < 2e-16 | < 2e-16 | < 2e-16 | 0.11941 | 0.5326 | - | - | - | - | - |
| **RaChr09** | < 2e-16 | < 2e-16 | < 2e-16 | < 2e-16 | < 2e-16 | < 2e-16 | < 2e-16 | < 2e-16 | 0.24147 | 0.29485 | 0.76432 | - | - | - | - |
| **RaChr11** | < 2e-16 | < 2e-16 | < 2e-16 | < 2e-16 | < 2e-16 | < 2e-16 | < 2e-16 | < 2e-16 | 0.02478 | 0.76158 | 0.74188 | 0.49727 | - | - | - |
| **RaChr12** | < 2e-16 | < 2e-16 | < 2e-16 | < 2e-16 | < 2e-16 | < 2e-16 | < 2e-16 | < 2e-16 | 0.26473 | 0.17016 | 0.62886 | 0.83086 | 0.29485 | - | - |
| **RaChr15(03ortho)** | < 2e-16 | < 2e-16 | < 2e-16 | < 2e-16 | < 2e-16 | < 2e-16 | < 2e-16 | < 2e-16 | 0.000058 | 0.138 | 0.0383 | 0.02014 | 0.06322 | 0.00393 | - |
| **RaChr15(14ortho)** | < 2e-16 | < 2e-16 | < 2e-16 | < 2e-16 | < 2e-16 | < 2e-16 | < 2e-16 | < 2e-16 | 0.02091 | 0.91637 | 0.62916 | 0.37411 | 0.83086 | 0.234 | 0.1282 |
